# Collective anti-predator escape manoeuvres through optimal attack and avoidance strategies

**DOI:** 10.1101/2024.03.26.586812

**Authors:** Palina Bartashevich, James E. Herbert-Read, Matthew J. Hansen, Félicie Dhellemmes, Paolo Domenici, Jens Krause, Pawel Romanczuk

**Affiliations:** Institute for Theoretical Biology, Department of Biology, Humboldt-Universität zu Berlin, Berlin, Germany; Research Cluster of Excellence “Science of Intelligence”, Technische Universität Berlin, Berlin, Germany; Faculty of Life Science, Humboldt-Universität zu Berlin, Berlin, Germany; Department of Fish Biology, Fisheries and Aquaculture, Leibniz Institute of Freshwater Ecology and Inland Fisheries, Berlin, Germany; Department of Zoology, University of Cambridge, Cambridge, UK; Aquatic Ecology Unit, Department of Biology, University of Lund, Lund, Sweden; IBF-CNR, Consiglio Nazionale delle Ricerche, Area di Ricerca San Cataldo, Via G. Moruzzi No. 1, Pisa 56124, Italy

**Keywords:** predator-prey interactions, collective response, fountain effect, agent-based model

## Abstract

The collective dynamics of self-organised systems emerge from the decision rules agents use to respond to each other and to external forces. This is evident in groups of animals under attack from predators, where understanding collective escape patterns requires evaluating the risks and rewards associated with particular social rules, prey escape behaviour, and predator attack strategies. Here, we find that the emergence of the ‘fountain effect’, a common collective pattern observed when animal groups evade predators, is the outcome of rules designed to maximise individual survival chances given predator hunting decisions. Using drone-based empirical observations of schooling sardine prey (*Sardinops sagax caerulea*) attacked by striped marlin (*Kajikia audax*), we first find the majority of attacks produce fountain effects, with the dynamics of these escapes dependent on the predator’s attack direction. Then, using a spatially-explicit agent-based model of predator-prey dynamics, we show that fountain manoeuvres can emerge from combining an optimal individual prey escape angle with social interactions. The escape rule appears to prioritise maximising the distance to the predator and creates conflict in the effectiveness of predators’ attacks and the prey’s avoidance, explaining the empirically observed predators’ attack strategies and the fountain evasions produced by prey. Overall, we identify the proximate and ultimate explanations for fountain effects and more generally highlight that the collective patterns of self-organised predatory-prey systems can be understood by considering both social escape rules and attack strategies.

## Introduction

High risk and high uncertainty situations require fast and robust reactions by individuals. In such scenarios, it is beneficial for individuals to rely on simple decision heuristics (1– 3). In groups, the interplay of these individual responses spreading through social interactions, and the influence of external environmental driving factors, results in the emergence of complex self-organised collective response patterns with non-trivial consequences for collective and individual performance. This is clearly manifested in the context of predatorprey dynamics in groups. For example, collective patterns such as “vacuoles”, “splits”, and “waves” are often produced when predators attack grouping prey (4–7). These collective patterns are thought to be the outcome of prey attempting to flee in ways that maximise their survival chances, and predators attempting to attack groups in ways that facilitate their hunting success. Surprisingly little is known, however, about why particular social escape rules are adopted by grouping prey to escape predators, the collective escape patterns that are produced by those rules, and whether these rules are robust to different attack scenarios.

One collective anti-predator response is the so-called “fountain effect”, whereby prey break into two subgroups, turning in an arched trajectory around an attacking predator and rejoining at its tail, visually appearing like a fountain (see Fig. 1A). This collective response appears to allow slower-moving prey to outmanoeuvre faster but less manoeuvrable predators (8, 9), while also allowing the separated subgroups to rejoin after the attack, retaining the benefits of belonging to a larger group. Fountain effects are currently understood to be a by-product of individual escape manoeuvres, where animals attempt to flee while maintaining the predator inside their visual fields, inducing angles of escape that depend on the predator’s position (10, 11). Similar to other escape rules of solitary prey (12–15), these models suggest prey can maximise their distances from the predator, but do not consider the role of social interactions during these escape manoeuvres. Indeed, optimal escape rules are likely to be affected by social interactions, given empirical studies show grouping prey often flee with more uniform escape trajectories than solitary ones (16). No models, however, have considered how groups should optimise their escape trajectories to escape from the predator while interacting with others during an attack.

**Fig. 1.**
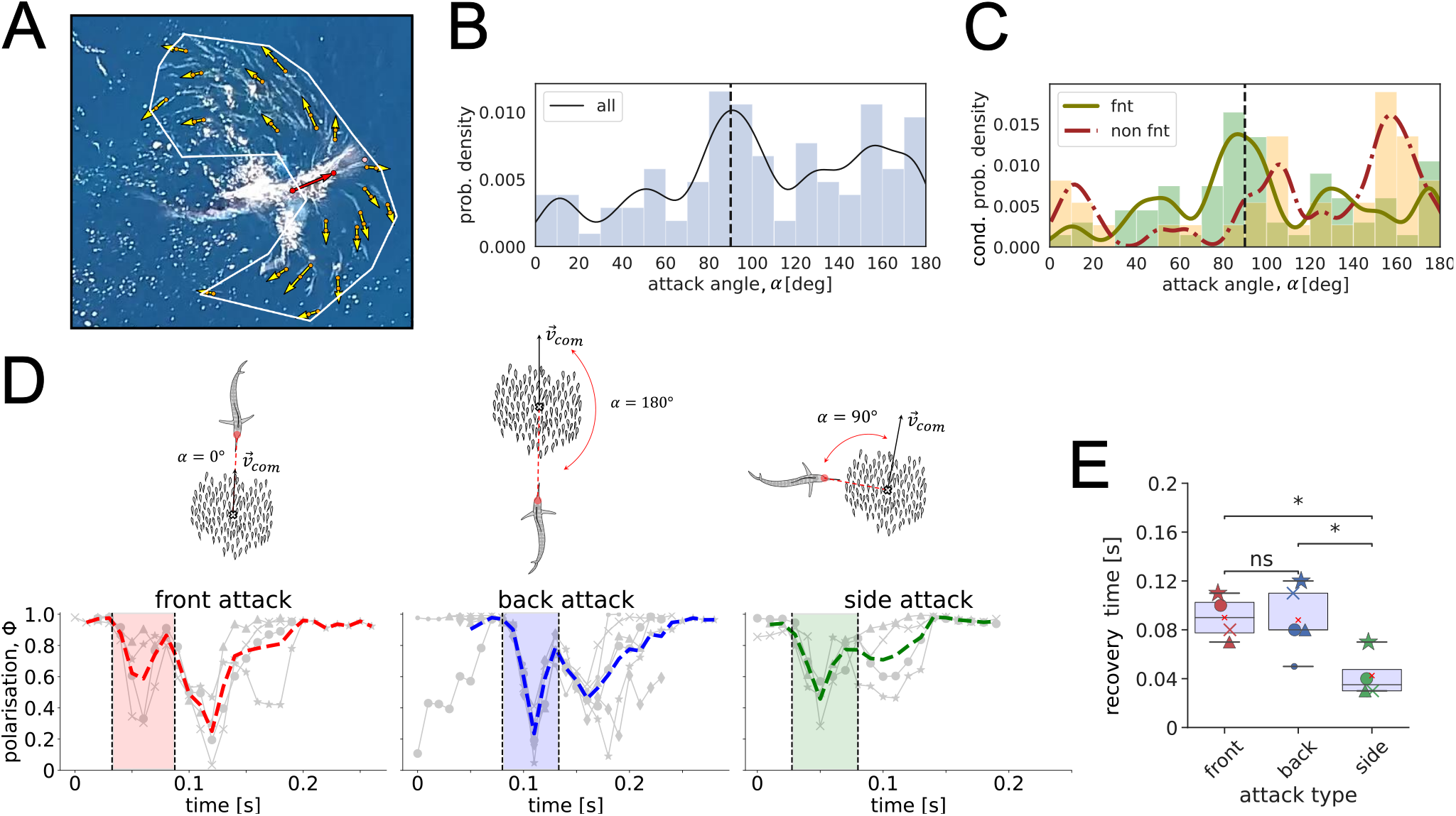
“Fountain effect” in the wild. **(A)** An instance of an annotated snapshot from the aerial footage over an open ocean, capturing the “fountain effect” performed by a school of sardines attacked by a striped marlin. Yellow and red vectors (defined by two dots as the head and the ‘tail’ on the fish) depict prey and predator velocity orientations, respectively. The arrows’ length is independent of the speed. The white polygon describes the borders of the school. The pink dot indicates the predator’s bill tip. **(B)** Probability density distribution of the predator attack directions α estimated between the predator’s velocity orientation and the heading direction of the prey’s school centre of mass over *n* = 104 recorded attacks. **(C)** Conditional probability density distribution of the predator attack directions based on the pattern of the observed anti-predator response, “fountain” (in green) or “non fountain” (in red). Each distribution is independently normalised according to the respective number of events (“fnt”: *n* = 67 and “non fnt”: *n* = 37), resulting in the rescaling of the bar heights (y-axis) compared to the entire distribution in **(B)**. Histograms’ overlap results in a third colour other than green (for “fnt”) and orange (for “non fnt”). **(D)** Prey polarisation Φ over time, estimated with 20 randomly annotated sardines as shown in **(A)**. Polarisation during each encounter is indicated by a grey line, while the thick line in colour shows the averaged Φ over the encounters of the respective attack type (14 attacks by 11 marlins). The shaded areas indicate the fountain window times (i.e., “start” and “end” times) averaged over each single encounter (grey line) and therefore not necessarily aligning with the peaks of the averaged polarisation (thick coloured dashed line). Attack types are schematically illustrated on top of **(D). (E)** Comparison of the collective prey recovery time τ between front, back, and side attacks (13 attacks by 10 marlins). Statistically significant difference among the attack directions is stated by the Kruskal-Wallis test followed by the Tukey test for multiple pairwise comparisons marked with *p* < 0.05 (*), ns = non-significant.

Fountain effects may also be the outcome of predators attempting to attack groups in ways that improve the likelihood of breaking groups apart or catching prey. Indeed, fountain effects are thought to occur when predators attack the centroid of groups from behind (17), as such attack directions may be a mechanism by which predators can break apart groups, improving capture success (18). However, in the wild (4, 19, 20) as well as in the laboratory (21), predators often target groups from other angles (e.g., from the side or from the front of groups). If the direction of an attack (in relation to a prey group’s orientation) impacts the pattern of the escape responses produced (22) or the distance predators can approach individual prey, predators should target groups from particular directions. In response, prey should adopt robust escape rules that maximise their survival chances across a range of attack scenarios.

Here, we investigate the mechanism and function behind the fountain effect in light of the conflicting interests of predators and prey. We do this by analysing aerial drone footage of striped marlin (*Kajikia audax*) attacking schools of sardines (*Sardinops sagax caerulea*) in the open ocean, quantifying fountain escape patterns and their dynamics following attacks on the groups from different directions. To understand the risks and rewards of particular escape rules and attack strategies, we extend a generic, spatially-explicit agent-based model of predator-prey interactions based on stochastic differential equations (23) to reproduce the fountain effect. Despite its common occurrence in nature, the fountain effect has rarely been captured in simulation studies (24–27), often involving hard-coded constituting elements or the absence of numerical analysis preventing an objective comparison with empirical data. Our model not only qualitatively captures properties of the fountain evasion, but also explains the advantages of using particular social escape rules. The escape rules used trade off maximising the distance of individual prey from the predator while minimising the time taken to rejoin the school after an attack. Moreover, the model makes predictions about why predators are observed attacking groups from particular directions. Overall, we highlight how fountain effects are the outcome of how predators or prey are attempting to attack or avoid capture in ways that benefit their respective success, offering an explanation for the emergence of fountain effects during predator-prey conflicts.

## Results

### Empirical data analysis

We recorded aerial footage of wild predation events on a school of Pacific sardines by striped marlins, using unmanned aerial vehicles (DJI, Phantom 4 Pro V2.0) flying at 20 m altitude, 10-30 km offshore Baja California, Mexico. During 19 minutes of the video recording (at 30 fps), we observed *n* = 136 attacks launched by individual marlins (see SI Appendix, Fig. S1) from different directions with respect to the school’s orientation, dashing one at a time through the prey school of ∼ 100 individual sardines. In 104 out of these 136 attacks, we were able to estimate the directionality of the dash by measuring the attack angle *α* from the heading direction of the prey school’s centre of geometry to the predator’s head (Fig. 1D-top). We refer to the values of *α* closer to 0*°*, 90*°*, or 180*°* as attacks from the front, side, or back of the prey school, respectively. Attacks from the side of groups (*α* ∈ [90*°*, 100*°*]) were the most common, followed by attacks from the back (*α* ∈ [150*°*, 180*°*]), with attacks from the front of groups (*α* ∈ [0*°*, 20*°*]) being the least common (Fig. 1B).

We qualitatively categorise escape manoeuvres by prey schools as “fountain” or “non-fountain” evasions (see SI Appendix, Figs. S2-S4). Fountains were defined as the prey splitting into two subgroups that moved in opposite directions to the predator, before subsequently rejoining behind the predator while turning to face its direction. Fountains occurred in the majority of attacks (*n* = 67 fountains versus *n* = 37 “non-fountain” evasions). Other non-fountain evasions involved the group sharply turning away without splitting, or the group being attacked from the front without splitting and turning away (see SI Appendix, Fig. S3).

Unlike in previous laboratory studies (11, 21), we observed fountain effects predominantly when attacks came from the side of the school (Fig. 1C); the direction that predators were most likely to attack from (Fig. 1B). Non-fountain evasions were most likely to be occurring when the prey were attacked from behind (Fig. 1C); the second dominant attack direction (Fig. 1B). To understand why attacks from the side and from behind of prey groups may be the most dominant ones, we quantified the properties of fountain manoeuvres when groups were attacked from different directions.

### Self-organised dynamics during and post-attack

Using custom Matlab code, we extracted the escape angles of the prey and their relative positions to the passing predator before and during a fountain effect. We were only able to quantify fountain effects this way in *n* = 30 attack sequences, owing to the poor visibility of individual fish in some videos due to the sea conditions. To ensure compatibility with the computational model (see below), we restricted our quantitative analysis to instances where the predator was moving in a straight line through the school without sharp turns during the attack (*n* = 16). In all other cases, the high variability of the predator turning manoeuvres in terms of direction, magnitude and timing, together with a limited number of observations, prohibited a quantitative comparison with model simulations.

For each attack, we labelled the head and tail of 20 evenly distributed and randomly sampled sardines, as well as the head and dorsal fin of the attacking marlin (Fig. 1A), every third frame (videos filmed at 30 fps) of each sequence. The sequences contained several frames *before* and *after* the fountain effect occurred. We defined a “fountain” window *t*_*w*_ based on the relative position of the predator in the prey school. We defined the “fountain” start, 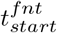, as 3 frames (0.1 s) before the predator’s bill is inside the prey group, defined by the polygon over the fish that are located along the school’s edges (Fig. 1A). We defined the “fountain” end, 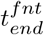, when the predator’s head (mouth) exited the prey school.

To analyse the directional organisation of individuals in the fountain over time, for each attack sequence, we quantified the degree of collective order through polarisation Φ as the mean of the velocity orientations (see Materials and Methods) of sampled sardines in each frame, for each attack direction separately (Fig. 1D). A significant decline in Φ indicates either fission or fusion events during and after the “fountain” window *t*_*w*_, respectively. Since the amount of annotated frames differed between the attack sequences, to align different attacks in time, we perform a time shift such that the minimum value of Φ (Fig. 1D) occurred at the same time between attacks. Note that despite 20 fish being sampled uniformly from the school because the school can split unequally (see SI Appendix, Fig. S6b), the average value of Φ can vary depending on how many individuals are on each of the split sides (on average, the fraction of annotations on split sides ranges in the proportion 35% − 65%).

The attacks from the back and from the front of the prey school were both described by the largest drop in polarisation compared to the attacks from the side. The largest decline in polarisation for the front attacks occurs on average after the predator leaves the school, highlighting that prey reverse the motion direction of the group after the predator has exited the school. That is, the prey school makes a 180-degree change in the orientation compared to the pre-attack. In the case of the back attacks, the decrease in polarisation is mainly observed during the fountain itself (i.e., when the predator is still inside the school), resulting from the prey splitting in opposite directions during the attack. The decline in Φ after the predator left was far less pronounced in the side attacks than in the front or back attacks. This is because for the side attacks, after the prey split, one of the group’s divided sides aligns with the other side without the two sides having directly opposing orientations towards each other (see SI Appendix, Fig. S6).

To quantify potential advantages gained for both predators and prey from attacks coming from certain angles, we quantified how long it took schools to reconfigure following an attack. Shorter recovery times may be advantageous to prey by reducing the likelihood of being re-attacked during a manoeuvre. Longer collective prey recovery times, on the other hand, could be advantageous to the predator by enhancing the disruption and fragmentation of prey groups, undermining their collective defences. To quantify the time it takes the prey school to reorganise after an attack, we propose the metric of *recovery time τ* (see Materials and Methods), which indicates the amount of time needed for the prey after an attack (i.e., after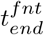) to get back to its initial pre-attack state (i.e., before 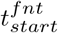). In the context of Fig. 1D, the latter corresponds to the highly aligned state, Φ →− 1. We find that the recovery time from a side attack is less than the recovery time of other attack directions (Fig. 1E), with the recovery from the front and back attacks taking similar amounts of time. Altogether, therefore, attacks from the side are most common, but the group can return to an aligned state more quickly when attacks come from this direction. Attacks from the back and front, on the other hand, see the largest changes to the group’s directional organisation and further take the group longer to return to an aligned state.

### Modelling the fountain effect

We use a spatially-explicit, stochastic agent-based model of predator avoidance adapted from (23). For simplicity, we restrict ourselves to a two-dimensional set-up. This simplification is empirically justified as the prey schools in our field observation are confined close to the water’s surface, and the prey evasion manoeuvres, and in particular the collective fountain responses, take place predominantly in a horizontal plane closely below the surface (see Fig. 1A). With our model, we explored what simple heuristics individual prey could use that ultimately lead to the collective dynamics observed in the empirical data above. In particular, we included a discrete escape manoeuvre into the avoidance force from the predator based on the concept of a *fleeing angle* Δ*α*_*flee*_ (Fig 2A). The response to the predator is modelled by a linear, distance-dependent repulsive force (“fleeing force”). For a vanishing fleeing angle Δ*α*_*flee*_ = 0*°*, the escape response occurs directly away from the predator along the vector 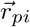 connecting the position of the predator and the prey agent *i*. For a finite fleeing angle, the fleeing direction is rotated by Δ*α*_*flee*_ in the direction away from the predator’s velocity vector 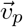 (Fig 2A).

**Fig. 2.**
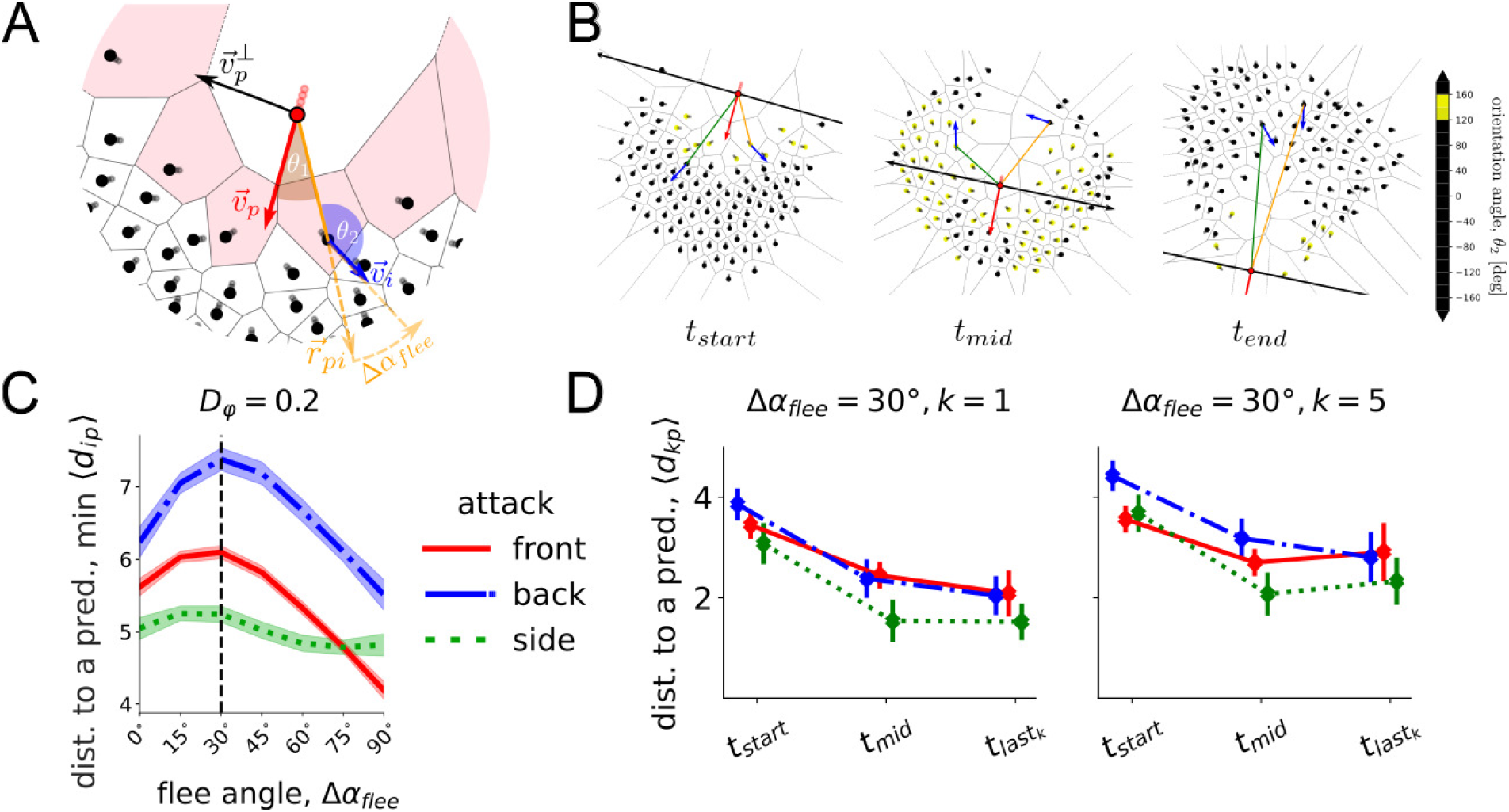
Optimal flee angle. **(A)** A snapshot from computational simulation modelling the “fountain effect”. The predator (red point) interacts with its first shell of Voronoi neighbours (black dots in pink cells), which are referred to as *direct responders*. All direct responders flee away with a repulsion force along the respective radius-vector 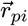 rotated on Δ*α*_*flee*_ in the direction away from the predator’s orientation 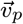. **(B)** The state of the simulation at the “start”, “middle”, and “end” of the “fountain”. These time points are defined by the relative position of the prey agents (black dots) to the black line, described by the vector 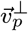. The predator moves straight with velocity 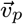 through the centre of mass of the prey school, which is reached by t_mid_. While flee angle as an “internal” behavioural parameter Δα_flee_ is set to 30*°*, due to the impact of social interactions the resulting orientation angle θ_2_ does not equal 150*°* for all individuals but occurs to be in the range θ_2_ ∈ [120*°*, 160*°*] for the prey agents coloured in shades of yellow. **(C)** The minimal distance to the predator min⟨d_i*p*_⟩ (averaged across all prey individuals i), which was ever achieved during the “fountain” window [t_start_, t_e*n*d_], depending on the flee angle Δα_flee_ with orientational noise intensity D_φ_ = 0.2 for simulated front, back, and side attacks. The curves are created by local regression with shading areas of 95 % confidence interval based on 40 simulation realisations for each attack type. The dashed vertical line shows the theoretical optimal flee angle 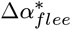 of a single escaping fish, as by Hall et al. (11). **(D)** Time-dependent distance changes between the predator and k closest prey. Distance to the predator ⟨d_*p*,k_⟩ averaged across its k nearest neighbours, using Δ*α*_*flee*_ = 30*°*, at the start, middle, and the end of the “fountain”. The end is specified by the time 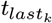 when there are only k prey agents left in front of the predator. The error bars indicate the standard error over 40 simulation realisations.

While directly escaping a predator, the prey agents align with other neighbours, although the strength of the flee *μ*_*flee*_ is set as the strongest among other social forces (see Materials and Methods). Note, given that the model also considers fluctuations in the heading of the prey with the intensity *D*_*φ*_, the flee angle Δ*α*_*flee*_ does not necessarily complement the resulting orientation angle *θ*_2_ of the prey. Namely, although we set the flee angle in the model, it defines the direction of the flee force, but not the final angle at which the agent flees. Since the fleeing force is additive to other social forces and orientational noise, despite being the strongest force among others, the final combination of forces defining the agent’s orientation *θ*_2_ is not necessarily *θ*_2_ = 180*°* − Δ*α*_*flee*_ (see Fig 2B with Δ*α*_*flee*_ = 30*°*). To ensure prey fusion after the split, we also replaced a distance regulating social force from (23) with a simple spring-like linear attraction-repulsion function (see Materials and Methods). As a result, we expanded the functionality of the original model (23), where the fountain effect had not been *a priori* observed. The model operates on the assumption that social forces modify only the prey’s orientation, keeping the speeds of both predator *v*_*p*_ and prey *v*_0_ constant (*v*_*p*_ = 2*v*_0_). The model speeds choice is consistent with the subsampled estimation of the predator-prey speed ratio (*v*_*p*_ : *v*_0_) in the empirical data (mean: 1.85, median: 1.88, range: [1.51, 2.14], standard deviation: 0.21) and does not show significant variations in the simulation results for the range *v*_*p*_ = [1.5*v*_0_, 2.0*v*_0_] (see SI Appendix for details, Fig. S8). Independent of the attack angle, we consider that the predator is directed towards the centre of the prey school and moving in a straight line. With this approach, we aim to relax the impact that variations in the predator trajectory have on the collective response of the prey, to comprehend its self-organised dynamics and to compare with the empirical data.

Unlike previous models used to generate fountain effects, e.g., (11, 27), we include social interactions between prey which are maintained during predator avoidance. In particular, focal individuals directly interact with the first shell of their Voronoi neighbours, whether these are other prey agents or the predator. Voronoi networks have been shown to approximate visual interactions in fish schools (28) and therefore provide a biologically realistic interaction neighbourhood. In the following, we refer to the first Voronoi shell of the predator’s neighbours as *direct responders* (see Fig 2A). That is, within the fleeing range *R*_*flee*_ these agents are affected by the flee force.

To allow comparison with the empirical data, we defined a “fountain” window *t*_*w*_ in the model based on the relative position of the prey to the orthogonal predator’s velocity vector 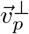. Namely, the vector 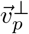 divides the space into the regions ‘ahead’ of and ‘behind’ the predator (Fig 2B). We define the “fountain” has started, 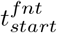, as soon as there is at least one prey individual ‘behind’ the predator while the rest of the group is ‘ahead’ of it. When the predator exits the school, we define the “fountain” end, 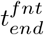 such that there is only one prey individual ‘ahead’, separated from the rest of the group which is left ‘behind’ (see Materials and Methods for details).

### Optimal flee angles and optimal angles of attack

We first tested the prediction that there is an optimal flee angle of the prey that maximises the distance of prey fish from the predator as it attacks (11), while socially interacting with others. To do this, we performed systematic numerical simulations of our model across different flee angles Δ*α*_*flee*_ (ranging from 0*°* to 90*°* with the step of 15*°*) for varying noise levels *D*_*φ*_ and different approach angles of the predator denoted as *front, back*, and *side* attacks, including their intermediate variations (see SI Appendix). We evaluated the optimality of the flee angle in terms of the mean distance of prey from the predator within the time window of the fountain effect. That is, the mean distance ⟨*d*_*ip*_⟩ of all prey *i* = 1, .., *N* to the predator is computed at each time step of *t*_*w*_, and the lowest mean distance of prey to the predator over the attack, noted as min ⟨*d*_*ip*_⟩, is selected. In this way, the minimal mean distance is strongly correlated with the maximal risk of predation within the school (18, 29, 30).

The model shows that regardless of the attack angle, the minimum distance from a predator peaks at around the same flee angle 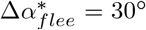(Fig. 2C). With this optimal flee angle, predators are more likely to get closer to prey when attacking from the side of groups, compared to the front and back of groups. This pattern is largely consistent regardless of the noise intensity of the flee angle (see SI Appendix, Fig. S7) and is consistent with previous theoretical results (10, 15, 31–33) for a single deterministic prey agent, i.e., in the absence of noise and interactions with the conspecifics. However, we note that here, the optimal flee angle 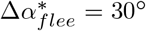 emerges in the absence of an explicit blind angle in prey as it was suggested based on theoretical consideration in Hall et al. (11), but within the context of the spatial self-organisation of the Voronoi interaction network. With an additional blind angle on top of the Voronoi neighbourhood, the optimal flee angle remains robust relative to the shift imposed by the blind angle, i.e., the resulting flee angle is the sum of the blind angle and the 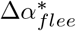 (see SI Appendix, Fig. S7).

We also evaluated the mean distance to the predator ⟨*d*_*p,k*_⟩ of only the *k* nearest neighbours at the start, middle, and end of the “fountain” (Fig. 2D). This allows us to assess at which moment of the fountain the predator gets closest to individual prey, and therefore the moment of greatest risk to individual prey. For the closest prey, i.e. when *k* = 1, the predator generally gets closest to the prey over 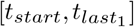 regardless of the approach angle *α*. As with the group metrics, with an optimal flee angle 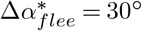, the back attack is more advantageous for the prey among other attack angles at the start *t*_*start*_, but the distance to prey levels off, so that by the end 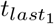 of the “fountain”, this distance is similar to that of front attacks. Side attacks, on the other hand, result in the lowest ⟨*d*_*p*,1_⟩ at all times compared to others, with no additional decline in 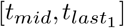.

For the *k* = 5 closest individuals to the predator (i.e., the average number of direct responders to a predator at a time), the distance ⟨*d*_*kp*_⟩ from the predator is initially larger for back attacks than side and front attacks at *t*_*start*_ and *t*_*mid*_. However, owing to an increase in ⟨*d*_*p*,5_⟩ towards the end of the fountain for front attacks, this distance ends up being equivalent to distances in back attacks by the end of the attack. Overall, side attacks generally result in the lowest distance to the predator for the closest *k* = 5 individuals and are consistent with the ones in Fig. 2C, indicating that the average minimal distance to the predator ⟨*r*_*ip*_⟩ over the group is correlated with the distance of a single (*k* = 1) individual. Our simulations, therefore, predict that predators will try to attack schools from the side as it allows them to get closer to prey *during* and *at the end* of the attack, during which the school performs the fountain manoeuvre.

### Impact of the flee angle on self-organised dynamics

To test whether our model incorporating an escape angle of Δ*α*_*flee*_ = 30^*°*^ captured the dynamics of the fountain effect observed in the empirical data, we computed the respective mean orientation angle of the prey fish (*θ*_2_) in relation to the fish’s position relative to the predator’s head (*θ*_1_) at each 10*°* in both the empirical data and model (Fig. 3 and see SI Appendix for details). This describes the average escape angles over 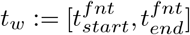 as the predator passes through and exits the school. Incorporation of Δ*α*_*flee*_ = 30*°* into the model qualitatively reproduced collective patterns of escape as observed in the empirical data.

**Fig. 3.**
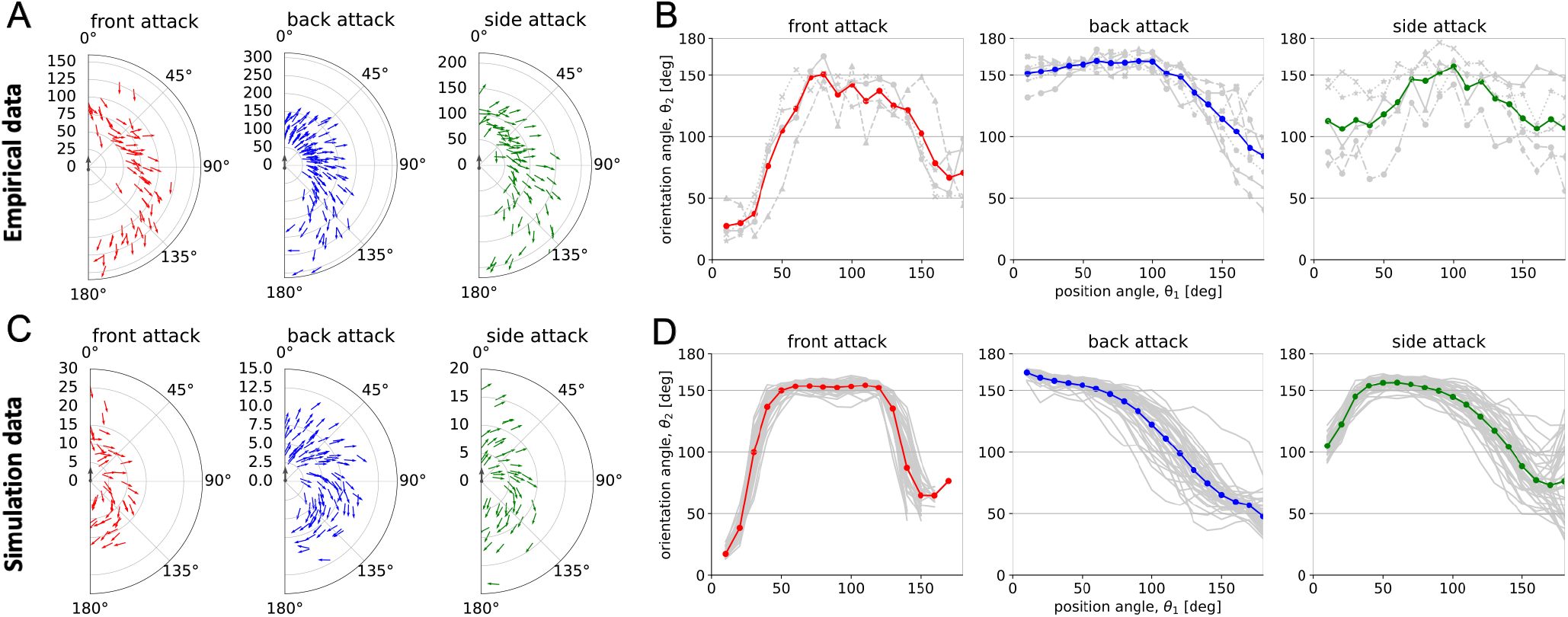
Self-organised dynamics of the fountain manoeuvre. **(A and C)** Polar plots illustrating aggregation of the attacks, where for each single attack the mean prey position and mean prey orientation in the increment of Δθ_1_ = 10*°* away from the predator’s head (placed in the origin) are computed over 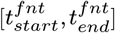. The amount of arrows in each sector Δθ_1_ is defined by the amount of respective attacks in **(A)** empirical and **(C)** simulation data with 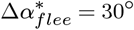. Radial y-axis indicates relative distance obtained from video data in pixel adjusted to the dimension of the manoeuvre.**(B and D)** Relationship between prey’s position angle relative to the predator (θ_1_) and prey’s orientation (swimming) angle (θ_2_) in **(B)** empirical and **(D)** simulation data with 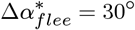. Each grey line represents a single fountain instance, while the thick line in colour shows the average over the instances. The results in **(D)** are based on 40 implementations, while in **(C)**, to prevent visual bias compared to the empirical data, we sampled an equivalent number of attacks as in **(A)**.

### Optimal post-attack recovery time

Although escaping from a predator at Δ*α*_*flee*_ = 30*°* increases the distance between the prey and the predator (Fig. 2C), such a relatively small flee angle can also result in longer arched trajectories (27). This, in turn, can lead to the two split subgroups taking longer time to recover (i.e., recoalesce) after being split (SI Appendix, Fig. S9). Shorter recovery times *τ* are likely to be beneficial to prey, as this allows them to exploit the benefits of larger groups, such as the dilution and confusion effects, sooner. Consequently, achieving shorter recovery times while retaining larger distances from the predator may be incompatible objectives for the prey. To assess how each flee angle Δ*α*_*flee*_ performs across these two objectives, we conducted a multi-objective analysis of our simulation results for different attack angles (Fig. 4A-C). For each attack direction, we find a set of fleeing angles Δ*α*_*flee*_ that creates a trade-off between the distance to the predator and the recovery time, forming the so-called Pareto-optimal solutions (see SI Appendix). There, improvements in one metric, i.e., increasing the distance from the predator, can lead to the worsening in the other, i.e., prolonging the recovery time. If we assume that there is no preference for one of the two aforementioned objectives (i.e., prey weight each objective with the same importance), the flee angles Δ*α*_*flee*_ that ideally balance recovery time *τ* with distance ⟨*r*_*i,p*_⟩ to the predator for each attack direction (front, back, side) are Δ*α*_*flee*_ ∈ *{*60*°*, 45*°*, 45*°}*, respectively (see SI Appendix, Table S1). However, the distance between prey and predator during an attack is a more immediate threat to life for the prey, and therefore this objective is likely to take stronger importance over decreased recovery time. If there is a strong preference for maintaining larger distances from the predator ⟨*r*_*i,p*_⟩ rather than on decreasing recovery times, we find that the respective solution on all Pareto fronts corresponds to Δ*α*_*flee*_ = 30*°*, regard-less of the attack angle (Fig. 4D). In this way, 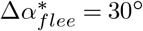 can be considered as an efficient heuristic, which prioritises maximisation of the distance from the predator. In other words, alternative Pareto-optimal solutions, different from Δ*α*_*flee*_ = 30*°*, enhance recovery time but tend to reduce the distance from the predator.

**Fig. 4.**
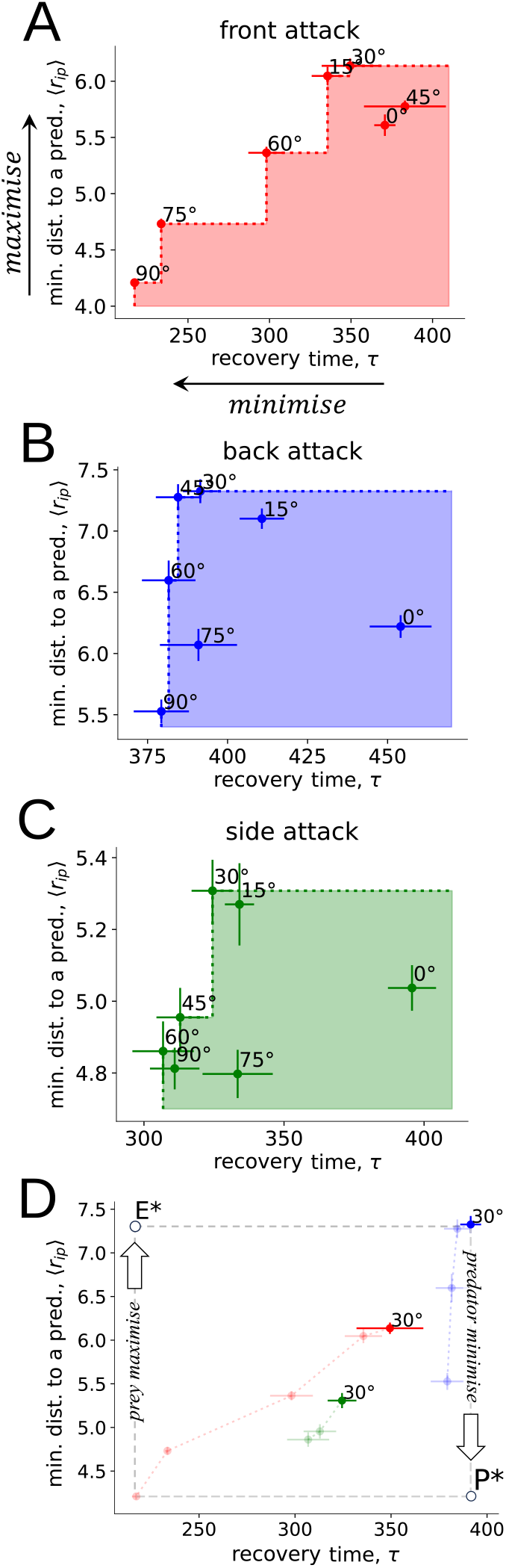
Conflict between collective recovery time and distance from the threat. **(A-C)** Pareto charts of flee angles Δ*α*_*flee*_ as prey movement strategies, with the utopia point as the maximum value of minimal averaged prey distance to a predator ⟨r_i,*p*_⟩ and the minimum value of collective recovery time τ (indicated by arrows in A). Each dot represents the mean value from 40 simulation implementations, and the horizontal and vertical bars indicate the standard errors for the corresponding metric. Nondominated solutions that form the Pareto front and reveal the trade-off lie on the chart’s “steps” covering the others (in shaded areas). **(D)** Perspective on the Pareto fronts of flee angles, conditioned on the prey escape, as predator’s attack strategies referred to front (in red), back (in blue), and side attacks (in green). P* denotes the ideal solution for the predator, and E* for the prey.

We can further analyse the Pareto-fronts of prey escape behaviour from the perspective of the predator’s attack strategies, referred to as front, back, and side attacks (Fig. 4D). Conditioned on the Pareto-optimality of the prey’s response, we find that the flee angle Δ*α*_*flee*_ = 30*°* creates a conflict in the effectiveness of the predator’s attack strategies and prey avoidance. Indeed, predators are the least effective at attacking prey groups from behind, as prey can get furthest away from the predator when they are attacked from this direction, despite a longer recovery time (increasing *τ*). Our model instead predicts that predators would be most effective if they attacked groups from the side (Fig. 4D), despite the fact that this attack direction results in the shortest recovery time *τ* for prey compared to other attack directions (Fig. 4D; consistent with our empirical evidence, Fig. 1E). While shorter recovery times are not favourable for the predator, side attacks remain closest to the ideal solution for the predator across bi-objective space (Fig. 4D; see also SI Appendix for more details, Fig. S10). Furthermore, although our results suggest back and front attacks are suboptimal for the predator with respect to the distance to the prey, predators receive a secondary benefit from front or back attacks because of the related increase in the prey’s recovery time, potentially leading to increased dispersion.

## Discussion

Using an empirically motivated agent-based model of predator-prey interactions, we show that by combining social interactions with a fleeing angle, groups of prey can produce “fountain effects”, with prey avoiding predators by splitting into two subgroups before rejoining together. By viewing prey escape behaviour from a multi-objective perspective, we find a range of flee angles that create a trade-off between the distance that prey can keep from the threat and the time needed to re-organise back into a cohesive, polarised group following an attack. A flee angle of 30*°*, however, represents a robust and optimal choice across a range of attack directions to maximise the prey’s distance from the predator, albeit resulting in longer recovery times for the group. These results highlight how attack and avoidance strategies give rise to collective escape patterns in contexts where prey have no place to hide and when predators aim to break up prey groups to disperse and isolate individuals (34, 35).

Predators often attempt to fragment prey schools, as this reduces school size, which can increase capture success (18, 29, 36). Our empirical data collected under natural conditions show that predators generally attacked schools from the side and from behind, with side attacks most frequently leading to “fountains”. Previous studies using artificial predators have associated fountain effects with attacks from the back of the schools (11). While fountains from this attack direction can occur, “non-fountain” evasions, where the entire school evaded the attack cohesively in one direction rather than splitting and re-joining (see SI Appendix, Fig. S2), predominantly occurred when groups were attacked from the back (Fig. 1C). As our empirical data show, prey attacked from the back are able to adjust their escape to avoid fragmentation altogether, preventing the predator from moving through the school. According to our modelling predictions, despite side attacks resulting in shorter recovery times for prey (as also observed empirically), they represent a better solution in the predator’s bi-objective space compared to front or back attacks. Attacking from the side not only enables the predator to get closest to the prey school, which is likely to enhance its capture success (e.g., (18, 29)), but also represents the best compromise between approaching the prey while increasing their recovery time. The latter can be viewed as a capitalisation of the prey’s response (i.e., producing a fountain effect) when predators attack from the side.

From the prey’s perspective, prey should prioritise maximising their distance from the predator over minimising recovery time, as being close to a predator represents an immediate threat to life. Under these assumptions, a flee angle of 30*°* was predicted to be the best escape heuristic (Fig. 4A-C). Critically, this 30*°* flee angle emerged in the absence of an explicit blind angle of the prey and maximised distance to the predator across all attack directions. Therefore, this simple heuristic appears to maximise individual survival success across a range of attack scenarios. On the other hand, if we assume that prey seek a trade-off between distance to a predator and recovery time, our model predicts the best compromise would be to flee away at an angle of 45*°* (SI Appendix, Table S1). This choice would be robust for scenarios when prey are attacked from behind and from the side, but not from the front of the school.

Our results suggest there is an ongoing arms race between predators and prey in the strategies they are using to improve their own success at the expense of the other. If prey use an optimal flee angle of 30*°*, predators are more effective in attacking from the side of the schools, while prey are more effective in escaping when attacked from behind. While prey are relatively more manoeuvrable than predators (8), and therefore should have more control over which direction they are attacked from, our empirical data show that attacks from the side and behind were the two most common attack directions (Fig. 1B), potentially highlighting a battle for the attack directions each party prefers. While attacking from the back or side benefits prey or predators respectively, these outcomes introduce indirect conflicting consequences arising from the existing trade-offs. In particular, back attacks provide an indirect, rather than the primary, advantage to the predator in increasing the recovery time of the prey if a fountain is produced (despite prey keeping larger distances from the predator), while side attacks allow groups to recover faster (despite predators getting closer to the prey).

In general, while significant research effort has gone into understanding arms-races between predators and prey from an evolutionary perspective, little effort has been made into understanding the role of complex spatio-temporal dynamics of collective prey responding to (collective) predators. Recently developed technological capabilities (combining drone technology and video tracking) allow researchers to obtain quantitative data on complex predator-prey interactions at high spatial and temporal resolutions (37). Nevertheless, interpreting these data remains a fundamental challenge. Our modelling approach provides a means to address the inherent difficulty of inferring specific behavioural rules like the “desired” fleeing angle Δ*α*_*flee*_ purely from trajectory data. In these systems, the observed movement of individuals is a product of individual movement decisions, social interactions and stochastic effects. Thus, any observed fleeing angle that can be extracted from individual escape trajectories may significantly deviate from any true internal Δ*α*_*flee*_ rule. Here, modelling approaches combining evolutionary dynamics with spatially-explicit multi-agent dynamics are promising. They allow for a complementary investigation of the intricate interplay of spatio-temporal self-organisation and function, and on the emergence of specific spatio-temporal collective response patterns as evolutionary stable strategies (ESS) (23). According to (5, 23), the combination of high alignment strength with a strong flee strength (like in our model) implies that the “fountain effect” can correspond to an ESS. While further work is needed to confirm this, it is plausible to hypothesise its validity based on the ecological context. Nevertheless, an understanding of the rules underlying these collective escape patterns, and in particular the interplay between prey avoidance rules and predator attack angles, is relevant to a variety of predators that attack grouping prey. Furthermore, this approach will inform research about how groups of humans or autonomous vehicles can avoid threat and improve coordination (27, 38, 39).

Overall, our modelling predictions suggest that empirically prevalent attack directions can be to the advantage of either predator or prey. This indicates an ongoing arms race, where both predators and prey continuously adapt their tactics to enhance their own success relative to the other. As predators demonstrate effectiveness in attacking the groups from the side, whereas prey show agility and effectiveness in evading attacks from behind.

## Materials and Methods

### Agent-based model description

The prey agents are modelled as active Brownian agents *i* = 1, …, *N* with constant speed *v*_*i*_ = *v*_0_ and an angular noise *D*_*φ*_ based on the following stochastic differential equations of motion (23):

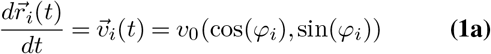

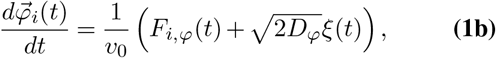

where 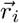 is the position vector of an agent *i*, ·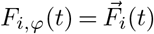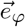. is a social force which is projected on the turning direction 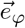 of an agent *i* and *ξ*(*t*) is a Gaussian white noise. The social force is composed as follows: 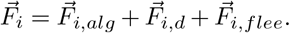 The agents align to each other with 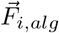 to keep the same direction of motion, they repel from each other with 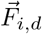 at short ranges (if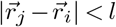), and attract to each other at long ranges (if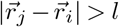) to stay cohesive:

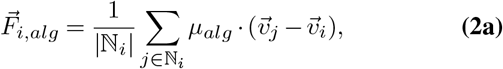

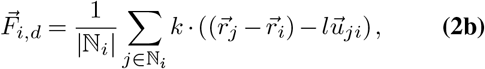

where 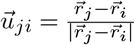 The neighbourhood subset of an agent *i* is denoted by ℕ_*i*_ and defined by the first Voronoi shell of neighbours. If a predator *p* is a neighbour of an agent *i, p* ∈ ℕ_*i*_, and 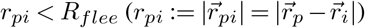 the agent *i* flees away from a predator according to

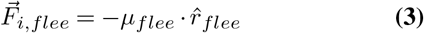

with 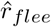 being the fleeing direction unit vector, which is given by the direction directly away from the predator 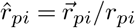 rotated by the additional fleeing angle Δ*α*_*flee*_ towards the rear of the predator:

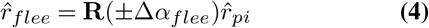

with **R**(*α*) being the rotation matrix by an angle *α* in 2D. The turning direction, i.e., the sign of the angle *±*Δ*α*_*flee*_, is always directed towards the rear of the predator, corresponding to the opposite direction to the predators velocity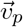 The predator moves with a fixed speed 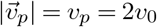.

The Euler-Maruyama method (40) is used to simulate the presented above stochastic differential equations with the time-step *dt*.

The predator is initialised relative to the prey’s group centre of mass 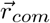 randomly within one of the defined angular regions (see SI Appendix) at the distance of *R*_*p*_ from the maximum remote prey agent from 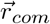 in this region. This initial relative positioning defines the predator’s *attack angle*. The right-hand or the left-hand positioning of the predator to the prey group is assigned randomly. The predator moves linearly to the updated position of the prey’s group centroid at each simulation step without noise in its velocity orientation 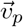 and continues moving straight after passing 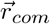. Before the predator is initialised, the prey group evolves for *T*_*eq*_ time units.

### Quantification of collective dynamics

For the empirical data, to estimate the degree of the collective order of the fish school before, during and after the escape, we computed polarisation Φ as the absolute value of the mean of the fish heading directions 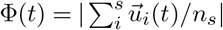, where *n*_*s*_ is the number of annotated fish in the frame *t* and 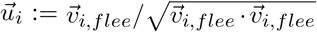 with 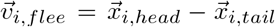, where 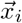 is the annotated location of the fish *i* head’s/tail’s coordinates (in pixels) in the frame.

### The start and the end of the fountain manoeuvre

Before the predator launches the attack, all prey agents *i* = 1..*N* are positioned in front of the predator: 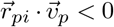. We defined the start of the “fountain effect” 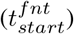 as soon as at least one prey individual *i* passes from frontal to rear relative position with respect to the predator, i.e., 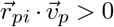. Accordingly, we defined the end of the “fountain effect” 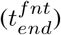 as soon as 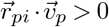 is true for all *i* = 1..*N*. These criteria also apply to empirical data, where 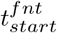 is defined by checking whether the annotated predator’s bill tip is within a polygon defining the borders of the prey school (see Fig 1A) using the Python module *mplPath*.

**Table 1.**
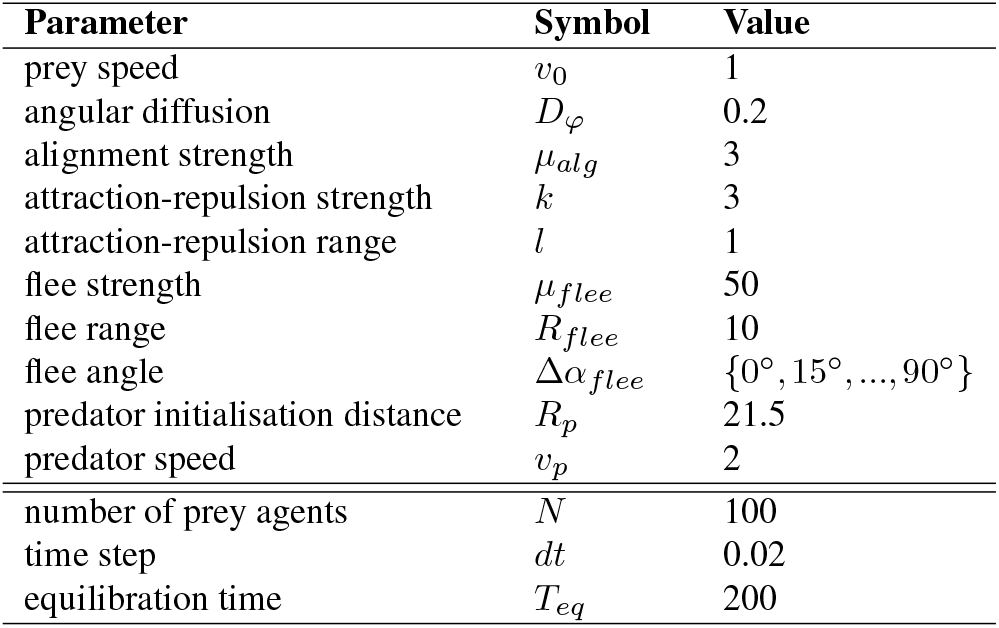
Model parameters. The values represent dimension-less units.

| Parameter | Symbol | Value |
| --- | --- | --- |
| prey speed | $v_0$ | 1 |
| angular diffusion | $D_\varphi$ | 0.2 |
| alignment strength | $\mu_{alg}$ | 3 |
| attraction-repulsion strength | $k$ | 3 |
| attraction-repulsion range | $l$ | 1 |
| flee strength | $\mu_{flee}$ | 50 |
| flee range | $R_{flee}$ | 10 |
| flee angle | $\Delta\alpha_{flee}$ | $\{0^\circ, 15^\circ, \dots, 90^\circ\}$ |
| predator initialisation distance | $R_p$ | 21.5 |
| predator speed | $v_p$ | 2 |
| number of prey agents | $N$ | 100 |
| time step | $dt$ | 0.02 |
| equilibration time | $T_{eq}$ | 200 |

### Collective recovery time

We computed the time-averaged mean polarisation 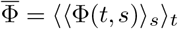 of the prey group over *n*_*s*_ = 40 simulations conducted in the absence of disturbance (predator) as a baseline order state of the simulated school. The respective standard deviation 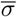 was computed over the concatanated array containing all polarisation values Φ(*t, s*) of the undisturbed prey at time *t* and each simulation *s*. In the simulations with a predator, we categorised the values of Φ(*t*) below the lower control limits, denoted as 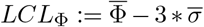, as indicative of a *perturbed* state of the prey group, and Φ(*t*) ≥ *LCL*_Φ_ as of a *recovered* state of the prey. To enhance the temporal stability of the data and minimise the impact of fluctuations, we applied the Savitzky-Golay filter on each time step of the time series difference *ŷ* (*t*) = *LCL*_Φ_ − Φ(*t*). As a result, based on the smoothed signal *ŷ* (*t*), the prey recovery time *τ* is the time point when *ŷ* (*t*) *>* 0 transitions to *ŷ* (*τ*) = 0.

For the empirical data, the lower control limits *LCL*_Φ_ are based on the mean 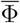 and 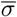 computed over the polarisation values 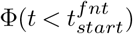 before the start of the fountain manoeuvre, satisfying Φ(*t, s*) *>* 0.8 to elimintate that the school can be still recovering from the previous attack and be in a perturbed state. The recovery time *τ* is set as a time point when polarisation values after the attack 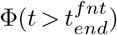 satisfy Φ(*t*) ≥ *LCL*_Φ_ such that the standard deviation of the polarisation sequence Φ(*t* ≥ *τ*) is close enough (*ϵ <* 0.01) to the standard deviation *σ* of the polarisation sequence 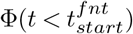 before the attack.

## ACKNOWLEDGEMENTS

PB, JK, and PR acknowledge funding by the Deutsche Forschungsgemeinschaft (DFG, German Research Foundation) under Germany’s Excellence Strategy – EXC 2002/1 “Science of Intelligence” – project number 390523135. MJH acknowledges funding by the Deutsche Forschungsgemeinschaft (DFG, German Research Foundation) grant - GZ:HA 9403/1-1 AOBJ: 67584. JEHR was supported by the Whitten Programme in Marine Biology, the Swedish Research Council (2018-04076), and the Office of Naval Research Global (N62909-21-1-2005). Special thanks are due to Max Licht for schematic illustrations of marlin and prey in Fig. 1D; to Korbinian Pacher for assistance in annotating the empirical observations; to Felipe Galván-Magaña at CICIMAR, La Paz, Mexico, and Magdalena Bay Whale Tours for assistance obtaining videos of striped marlin in the field.

## AUTHOR CONTRIBUTIONS

P.B. and P.R. designed the study; P.B. performed research; M.J.H., F.D., and J.K. acquired the empirical data; P.B. performed the numerical simulations and analysed the results with input from J.E.H-R., P.D., and P.R.; P.B. and J.E.H-R. wrote the paper. All authors commented on the manuscript.

## Supplementary Material

### Categorising “fountain” and “non-fountain” escape manoeuvres

Qualitatively, we relate “non-fountain” cases (see Fig. S3) to the attack events when the prey school is performing a sharp turn to the predator’s tail, but the whole school sticks together and does not split into two subgroups by either side of the predator unlike in the case of the “fountain effect” (see Fig. S2). We also quantified the shape of the prey group based on the points of the annotated polygon over the fish positioned on the edges of the school as the ratio of its convex hull perimeter to the polygon’s perimeter to define the level of the group’s convexity *C* during the attack (Fig. S5). Fig. S4 shows the convexity measures of the prey group’s shapes for “fountain” and “non-fountain” evasions in the middle (*C*_*m*_) and at the end (*C*_*m*_) of an attack. “Fountain” evasions are characterised by lower measures of convexity compared to “non-fountain” ones at the end of attacks, i.e. *C*_*e*_ *<* 0.9, which indicates a split in the group, confirming our qualitative classification. However, there are some borderline cases (typical for the front attacks) in the case of the “fountain” evasions, where the annotated group’s shapes do not undergo significant convexity changes, as prey are splitting in a more streamlined manner around the predator (see for instance, Fig. S2c,d). However, these are distinguishable through observation as the individual prey (more than 2 individual fish) are evidently located by both sides of the predator (Fig. S2d) compared to the “non-fountain” cases (Fig. S2f).

### Individual predator identities

To ascertain individual predator identities between attacks, we followed individual predators until they swam outside the video frame and could no longer be identified. This way, the predators labelled with the same ID are the same predator making multiple different attacks. However, we can not be absolutely certain about different predator IDs as the predators frequently left the frame of the video, such that it is likely that a subset of the attacks were made by the same predators at different times. Fig. 1B (in the main text) contains *n*=104 attacks performed by *N* = 55 (potentially) unique predator IDs. Among them, there are *N* = 32 unique predators who attacked only once, while the other *N* = 23 attacked at least twice. Out of these *N* = 23, *N* = 14 predators were identified to perform between at least *n* = 3 and *n* = 5 attacks at maximum (*N* = 6 predators did each *n* = 3 attacks, *N* = 4 predators did each *n* = 4, and another *N* = 4 predators did each *n* = 5). We subsampled *N* = 8 predators with *n* = 4 attacks each (see Fig. S1) and performed a repeated measures ANOVA to test if the individual predatory fish are behaving differently, as well as if there is a difference in the trial. There was no statistically significant difference in attack angles between individuals (*F*(7, 21) = 0.455, *p* = 0.855) and also no difference in the trial number (*F*(2.081, 14.57) = 1.491, *p* = 0.258).

Fig. 1D (in the main text) is based on *n* = 14 (4 from the front, 4 from the side, 6 from the back) attacks by *N* = 11 different predators. In one back attack the fish school was not able to recover before the next attack, therefore Fig. 1E based on Fig. 1D (in the main text) incorporates only *n* = 13 attacks. These attacks were launched by *N* = 10 different predators. Each attack type has no repeated attacks by the same predator (Front: “J” (star); “F” (circle); “E” (triangle); “G” (cross). Back: “J” (star); “C” (cross); “B” (circle); “I” (dot); “D” (triangle). Side: “E” (star); “A” (circle); “D” (triangle); “K” (cross)). Between the attack types, *N* = 3 predator individuals performed repeated attacks: “J” - front and back, “E” - front and side, “D” - back and side.

### Estimation of relative predator-prey speeds

An estimation of predator and prey speed was calculated by tracking the positions of both predator and prey using aerial video of attacks (N=13). These attacks were selected based on two criteria: (i) the drone was stationary (corroborated with the flight log) and, (ii) the position of the predator and an individual sardine were visible at each consecutive frame during a full second of the attack (i.e., 30 frames or 60 frames depending on the video). The frames were exported from each video using VirtualDub (41) and imported into ImageJ (42). A stereo camera (90 cm. in length), visible in each of the exported frames, was used as a scale object. Using the manual tracking plugin MTrackJ (43), the trajectory of the predator was measured by tracking the space between its pectoral fins at each frame. The trajectory of the sardine was measured by tracking the mid-point of its body for the same set of frames. The cumulative distance travelled by both predator and prey over every frame was calculated to get speed (cm/s): predator (mean *±* se = 334.76 *±* 11.74, range = [257.95, 397.75]); prey (mean *±* se = 183.33 *±* 8.05, range = [125.36, 228.62]). Predator speed was divided by prey speed to calculate the predator:prey speed ratio (mean *±* se = 1.85 *±* 0.06, range = [1.51, 2.14]).

### Model choice of relative predator-prey speeds

In our modelling approach, the assumed individual predator response is given by an evasion manoeuvre away from the predator, with an additional directional bias given by the fleeing angle towards the tail of the predator. This assumption is in line with previous studies (11, 21), and is also essential for the production of robust fountain-like collective responses as observed empirically. We should note that within a schooling manoeuvre, some of the early responses near the predator may correspond to what can be considered “escape responses” from a physiological perspective, as observed in previous work on schooling fishes (16, 22). These are typically fast accelerations away from a suddenly emerging threatening stimulus, with large directional variability in solitary fish (32, 44) and more uniform responses in schooling fish (16). For schools of pelagic fish being hunted by marine predators, such escape responses are likely to occur only near the predator as a last attempt to avoid capture but are unlikely to determine the overall schooling manoeuvres. Given their approximately 20 cm length (estimated mean *±* sd = 19.42 *±* 1.92 cm) and their speed of 1.83 m/s, prey were likely to swim near their maximum swimming speed and well beyond their maximum sustainable speed (estimated to be 1.9 and 0.6 m/s, respectively, based on a 20 cm long teleost (45)), with little capacity for further acceleration. This is also a main argument for considering for simplicity a constant speed of both predator and prey. Here, we note that in a previous modelling study, extending the model towards variable prey speed, and in particular including acceleration in the escape response, did not yield any qualitative changes in the collective escape dynamics (23).

### Polar plot illustration of fountain escape dynamics

For each attack encounter, 20 random fish were annotated each third frame up until the predator left the school. First, their positions and orientations were mapped into polar coordinates relative to the predator (the origin of the coordinate system), thereby aggregating all annotated frames of the particular attack at once. Secondly, the angular space was partitioned into 10° increments, with the average position and orientation of the fish within each region computed relative to the predator for this encounter. This procedure was repeated for each attack encounter separately. As a result, Fig. 3A in the main text illustrates the aggregation of all attacks of each type with mean prey position and mean prey orientation in the increment of 10° away from the predator. The amount of arrows in each sector is defined by the amount of respective attacks. To avoid visual bias in Fig. 3C in the main text, for the simulation data 20 agents were uniformly selected at random from the prey group at each iteration and the same number of attacks for each type as in the empirical data was used.

### Model sensitivity to predator attack direction, orientational noise, and prey blind angle

Our simulation results (the first row of Fig. S7) suggest that *side* and *front-side* attacks do not significantly differ from each other in terms of the closest distance attained to a predator ⟨*r*_*ip*_⟩, and the same holds for the *back-side* and the *back* attacks. This implies that one can consider *front-side* with *side* attacks as a one *side*-area attack category, and *back-side* with *back* as a one *back*-area attack category in the simulation. In the main text, we focus on the extreme attack angles, referred to as back, side, and front attacks initialised at 180°, 90° and 0° respectively with additive noise in the range (0*°*, 30*°*). Their relative performance is robust at various noise levels *D*_*φ*_ in the absence of an additional prey blind angle (the first row of Fig. S7). We find that *D*_*φ*_ = 0.2 is the optimal noise intensity of fluctuations in prey orientation, where the distance to the predator ⟨*r*_*ip*_⟩ is maximised in case of the back attack (i.e., facilitating prey escape) and minimised in case of the side attack (i.e., facilitating predator success). To note, *D*_*φ*_ = 0.2 remains optimal in terms of ⟨*r*_*ip*_⟩ regardless of the prey’s blind angle within its feasible range (i.e.,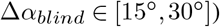). With an additional prey blind angle Δ*α*_*blind*_ on top of the Voronoi tessellation, the optimal flee angle 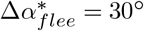 remains robust in case of the front attack and shifts to 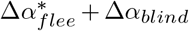 in case of back and side attacks for Δ*α*_*blind*_ ∈ [15*°*, 30*°*]) (see Fig. S7).

### Multi-criteria analysis of simulated prey escape

The bi-objective space (*f*_1_, *f*_2_) := (⟨*r*_*ip*_⟩, *τ*) of the minimal averaged prey distance from a predator ⟨*r*_*ip*_⟩ and collective prey recovery time *τ* is plotted to identify the Pareto-optimal prey fleeing angles and predator’s attack directions. Concerning the effectiveness of the prey escape, a flee angle 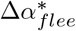 is Pareto optimal if it is not dominated by any other solution, i.e., there is no other Δ*α*_*flee*_ for which 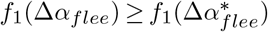 and 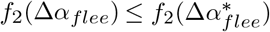, and there is a strict inequality for at least one objective. A set of all Pareto optimal solutions forms the Pareto front (PF). Fig. S10 contains the PFs of prey fleeing angles identified for each attack direction.

To identify the best compromise solution that comes as close as possible to achieving the best values for both objectives *f*_1_ and *f*_2_ simultaneously, we use the *ideal point* approach (46). For each PF (depending on the attack type), we construct the ideal point *I* from the respective PF’s *extreme solutions*, which correspond to the optimal solutions for the individual objective, such that 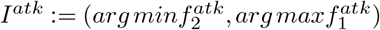) within the attack type. The best compromise solution is the one on the PF with the minimal Euclidean distance *ρ* from the respective ideal point in the normalised objective space (i.e., the values of *f* 1 and *f* 2 are in the range [0, 1]). Table S1 shows the Pareto optimal fleeing angles Δ*α*_*flee*_ for each attack type with respective Euclidean distance *ρ* from the ideal point *I*^*atk*^ of the corresponding attack type. That is, Δ*α*_*flee*_ = 45*°* is the best compromise solution for the prey in back and side attacks, while Δ*α*_*flee*_ = 60*°* is the best one for the prey in front attack.

By placing PFs of optimal prey escape angles for each attack type in the same objective space (Fig. S10), we can construct PFs of optimal attack direction from the perspective of prey and predator. We assume that the best for the prey would be to maximise the distance away from the predator (i.e., *max* ⟨*r*_*ip*_⟩) and minimise their recovery time (i.e., *min τ*), while the opposite is the best for the predator (i.e., *min* ⟨*r*_*ip*_⟩ and *max τ*). Based on the respective extreme solutions, we find the prey’s ideal 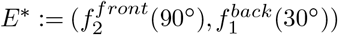 and the predator’s ideal 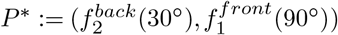 points (see Fig. S10). This way, the whole front-PF and the part of the back-PF (with Δ*α*_*flee*_ ∈ {30°, 45°, 60°}) correspond to Pareto optimal attack directions for the prey, while the whole side-PF and the whole back-PF are Pareto optimal for the predator. By computing the Euclidean distances *ρ*_*E*_*\** and *ρ*_*P*_ *\** from the prey’s *E*^*\**^ and predator’s *P* ^*\**^ ideals to the points on these PFs, we can estimate how certain attack directions, conditioned on the particular prey fleeing angle Δ*α*_*flee*_, balance the trade-off between distance from the predator and the recovery time, when both objectives (*f*_1_, *f*_2_) are taken into consideration. Front attack with Δ*α*_*flee*_ = 60*°* corresponds to the best attack direction for the prey (i.e., with the minimal *ρ*_*E*_*\**, Table S1). Back attack with Δ*α*_*flee*_ = 90*°* is the closest to the predator’s ideal point *P* ^*\**^, while side attack with Δ*α*_*flee*_ = 45*°* is the second-closest one. However, Δ*α*_*flee*_ = 90*°* is the extreme solution for the prey (i.e., with the minimal recovery time) without considering the second objective (i.e., distance from the predator) within the back attack (*ρ* = 1, Table S1). Therefore, conditioned on the prey response, side attack with Δ*α*_*flee*_ = 45*°* is the best attack direction for the predator (i.e., with the second smallest *ρ*_*P*_ *\**, Table S1).

**Table S1.** Euclidean distances from each Pareto front solution Δ*α*_*flee*_ to: (1) prey ideal point within attack type (*ρ*), (2) prey ideal point *E*^*\**^ across attack types (*ρE*^*\**^), and (3) predator ideal point *P* * across attack types (*ρP* *). The cross (x) denotes the solutions which are dominated (i.e., not Pareto optimal) with respect to the corresponding ideal point. The values of 1 indicate the extreme solutions on the Pareto front. The minimal value in each column corresponds to the optimal solution that is best balanced across both objectives (i.e., prey distance from the predator and prey recovery time after an attack).

| attack type | $\Delta\alpha_{flee}$ | $\rho$ | $\rho_{E^*}$ | $\rho_{P^*}$ |
| --- | --- | --- | --- | --- |
| back attack | 30° | 1 | 1 | 1 |
|  | 45° | 0.430 | 0.960 | 0.985 |
|  | 60° | 0.447 | 0.972 | 0.769 |
|  | 90° | 1 | x | 0.429 |
| front attack | 30° | 1 | 0.850 | x |
|  | 15° | 0.894 | 0.792 | x |
|  | 60° | 0.732 | 0.782 | x |
|  | 75° | 0.739 | 0.837 | x |
|  | 90° | 1 | 1 | 1 |
| side attack | 30° | 1 | x | 0.521 |
|  | 45° | 0.862 | x | 0.510 |
|  | 60° | 1 | x | 0.529 |

**Fig. S1.**
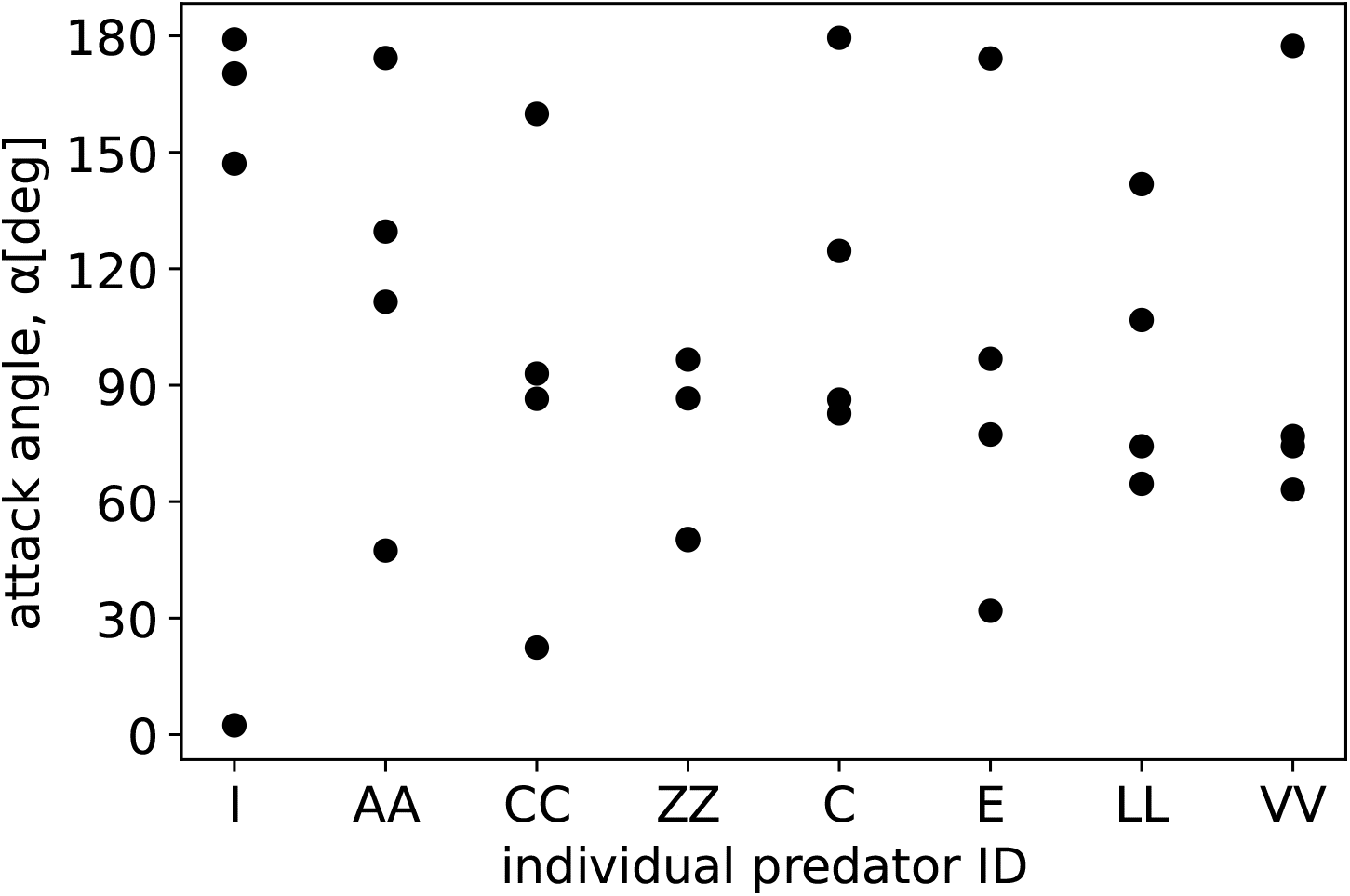
Each dot shows the attack angle by an individual marlin (a subsample of 8 individual marlins with 4 attacks each), estimated by an angle between the heading direction of the prey school’s centre of geometry to the predator’s bill tip 3 frames (0.1 s) before entering the school. According to the repeated measures ANOVA test, there is no statistically significant difference in attack angles between individuals.

**Fig. S2.**
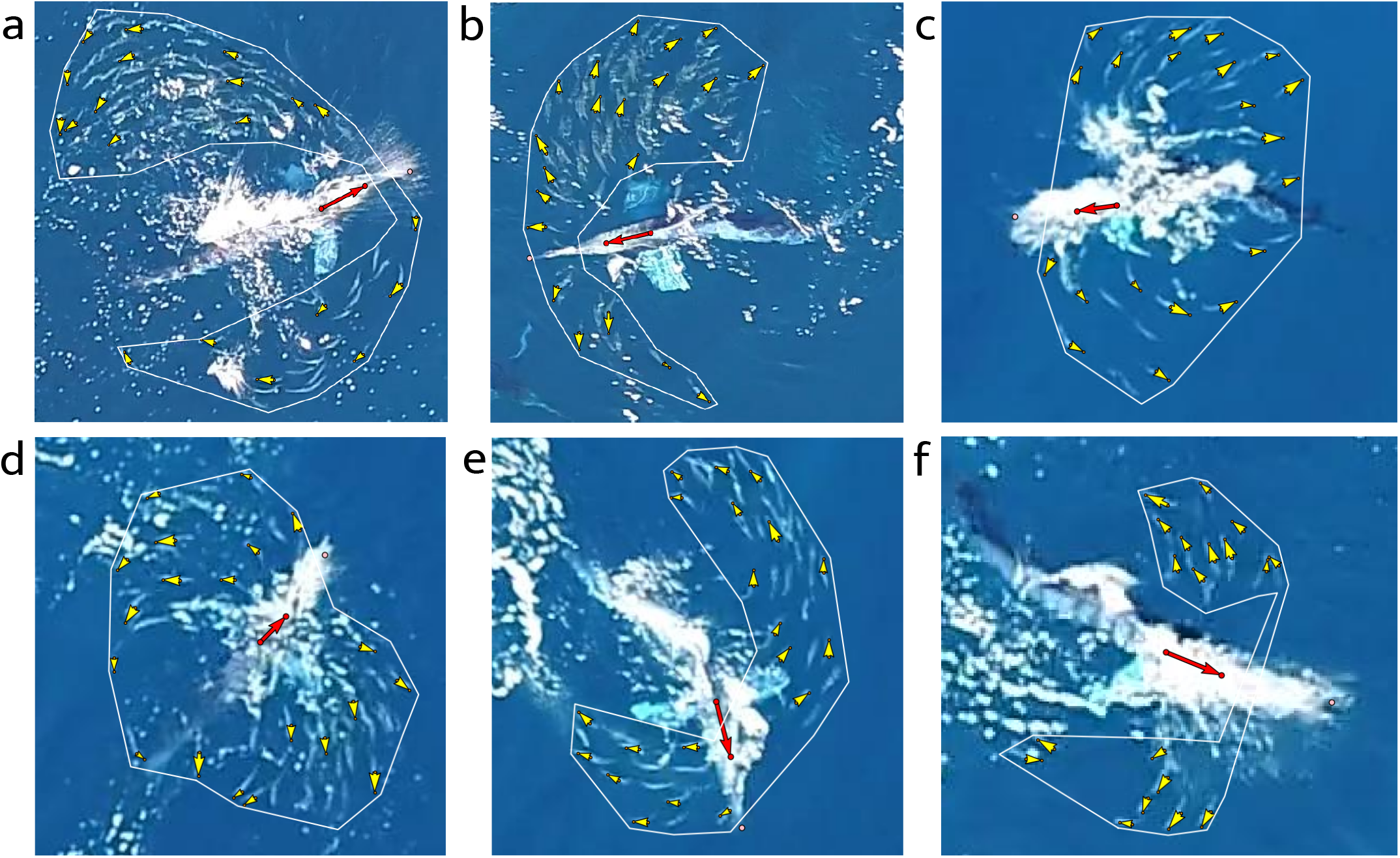
Instances of the zoomed aerial footage with the “fountain effect” during the attack by a striped marlin. The white polygon captures the borders of the prey school. Yellow arrows show the direction of motion of the annotated prey individuals, while the red arrow (connecting the marlin’s fin and the head) depicts the predator’s orientation. The arrows’ length is independent of the speed. The pink dot indicates the predator’s bill tip.

**Fig. S3.**
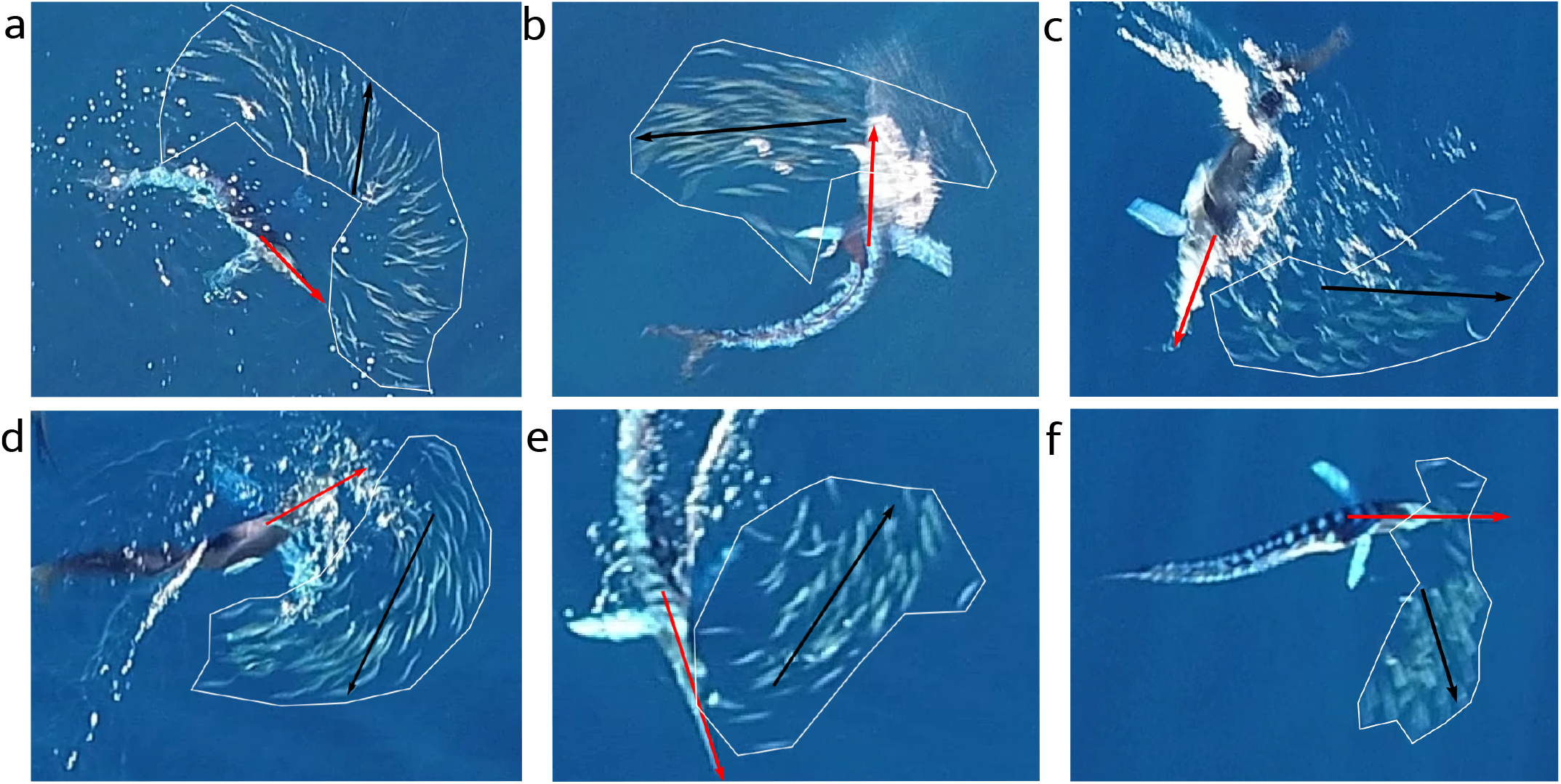
Instances of the zoomed aerial footage with non-fountain escape manoeuvres during the attack by a striped marlin. The white polygon captures the borders of the prey school. The red arrow (connecting the marlin’s fin and the bill tip) depicts the predator’s orientation. The black arrow inside the polygon shows the general direction of motion of the whole prey school. The arrows’ length is independent of the speed.

**Fig. S4.**
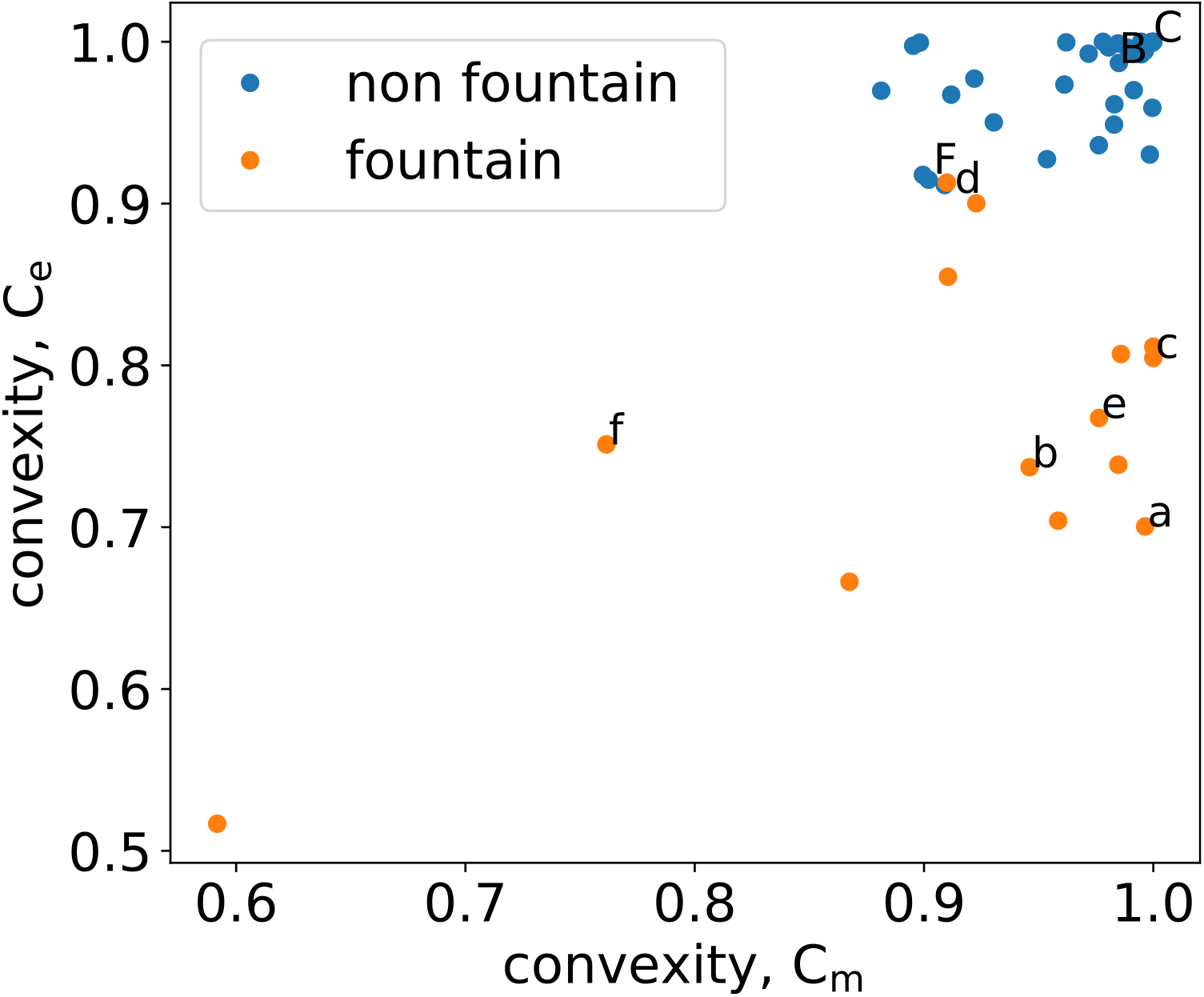
Each dot shows the convexity of the polygon defining the borders of the prey school geometry in the middle *C*_*m*_ and in the end *C*_*e*_ of an attack over annotated *n* = 37 non-fountain (in blue) and *n* = 14 fountain (in orange) evasions. The middle of an attack is defined as an approximate point in time between the start and the end of an attack. The start of the attack is defined as 3 frames before the predator’s bill is on one level with the prey fish, such that a drawn perpendicular line to the bill’s tip intersects at least with one sardine. The end is defined as when there is no more any fish in front of the mouth and the bill of the predator. The capital letters correspond to the non-fountain evasions in Fig. S3 and the lower case letters indicate the respective fountain evasions in Fig. S2.

**Fig. S5.**
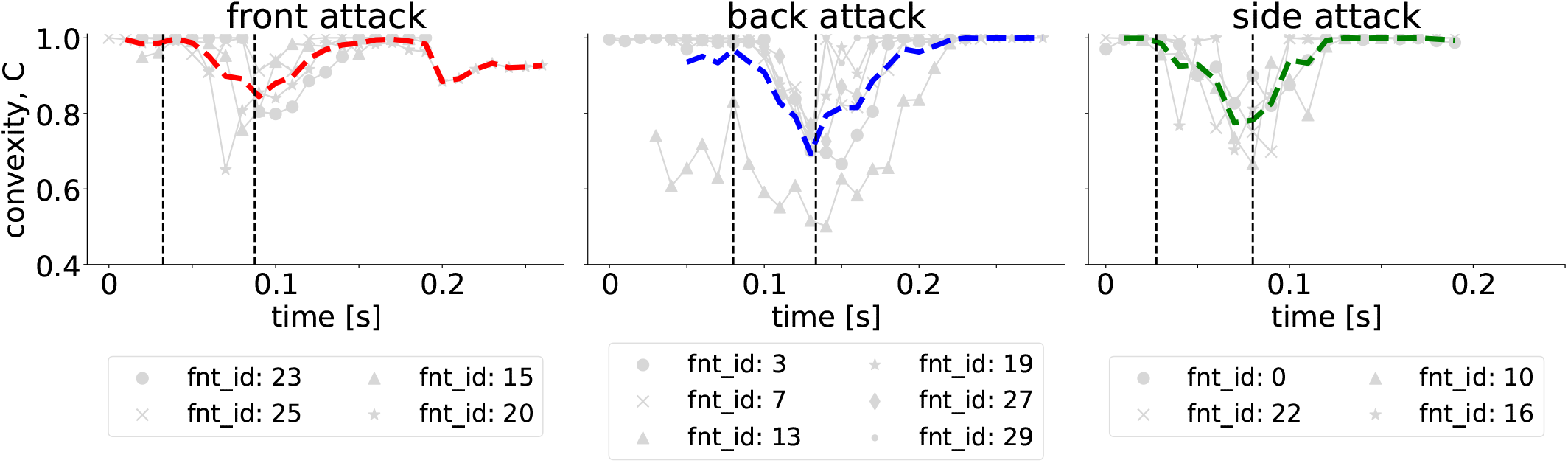
Convexity *C* of the annotated polygon defining borders of the prey school over time in case of the fountain manoeuvre. The dashed vertical lines mark the averaged “start” and “end” times of the “fountain” over the encounters. Convexity of the school during each encounter is indicated by a grey line, while the thick line in colour shows the averaged *C* over the encounters of the respective attack type. Each marker corresponds to a particular fountain manoeuvre (id) and is consistent with the markers in Fig. 1D in the main text.

**Fig. S6.**
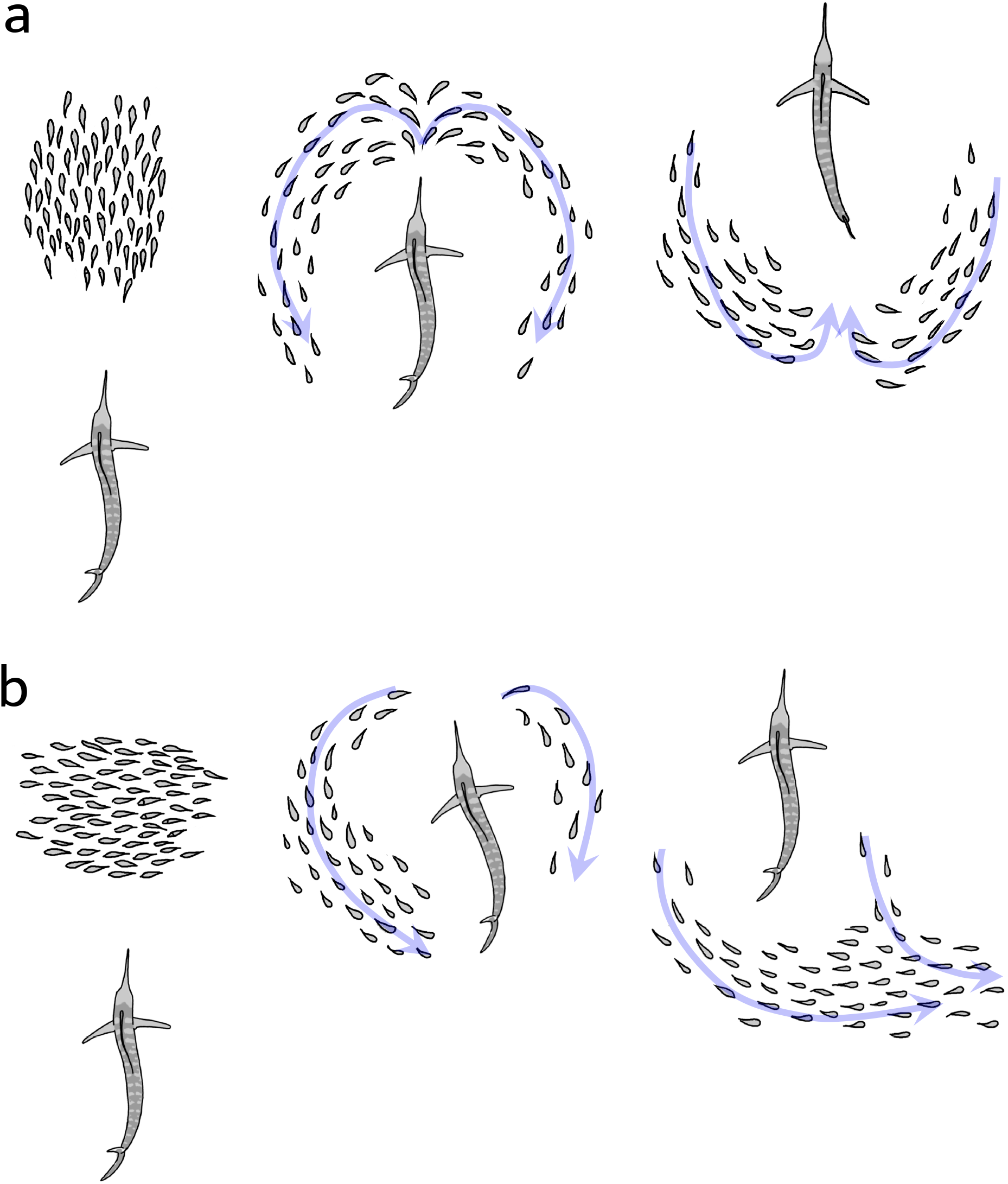
Schematic illustration of the prey group splitting and re-joining behind the predator during the fountain manoeuvre, when prey is attacked from behind (a) and from the side (b). In case of the back attack (a), prey subgroups move towards each other to rejoin and re-face the same direction of motion as the predator. In the case of the side attack (b), prey subgroups realign without moving in opposite directions towards each other. Blue arrows show prey subgroups’ general direction of motion during the split.

**Fig. S7.**
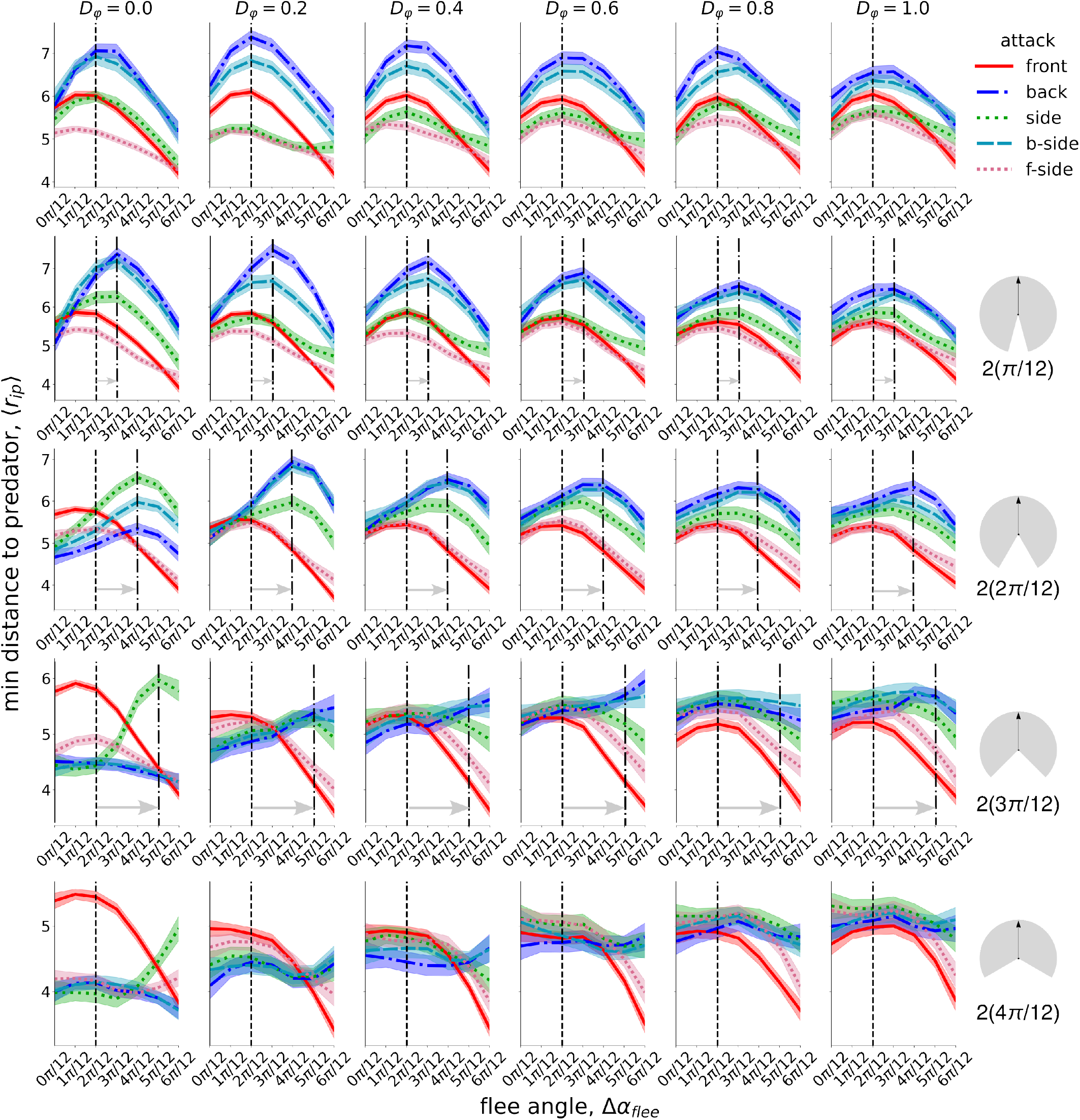
The minimum distance to the predator ⟨*r*_*ip*_⟩ := *min*⟨*d*_*ip*_⟩ (averaged across all prey individuals *i*) achieved during the “fountain” window, depending on the flee angle Δ*α*_*flee*_, prey orientational noise levels *D*_*φ*_, and 5 attack directions. The curves are created by local regression with shading areas of 95 % confidence interval based on 40 simulation realisations for each Δ*α*_*flee*_. The dashed vertical line shows the theoretical optimal flee angle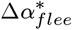, as by Hall et al. (11). The grey horizontal arrow indicates the shift from 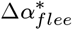 on the prey blind angle depicted by the empty sector in a grey circle on the right-hand side of each row. The first row states the results without a blind angle over the Voronoi interaction network. In teleost species, a blind zone of 10*°* − 30*°* to the rear on either side of the fish has been reported (11, 47). The predator:prey speed ratio is set to 2 : 1.

**Fig. S8.**
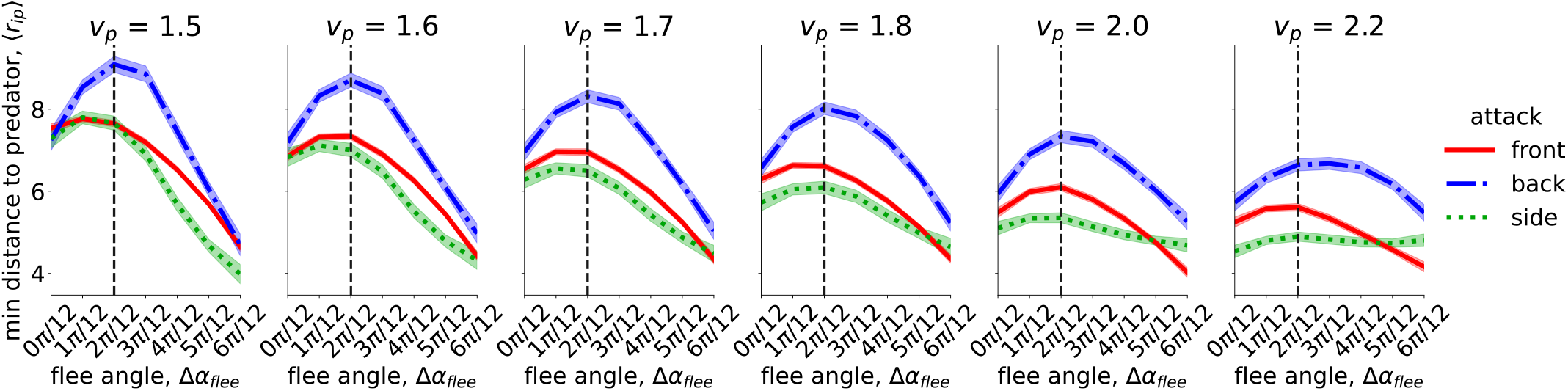
The minimum distance to the predator ⟨*r*_*ip*_⟩ := *min*⟨*d*_*ip*_⟩ (averaged across all prey individuals *i*) achieved during the “fountain” window, depending on the flee angle Δ*α*_*flee*_ and the predator:prey (*v*_*p*_ : *v*_0_) speed ratio with *v*_0_ := 1. The value of the prey angular noise is set to *D*_*φ*_ = 0.2 and there is no additional blind angle over the Voronoi interaction network. The curves are created by local regression with shading areas of 95 % confidence interval based on 40 simulation realisations for each Δ*α*_*flee*_ and *v*_*p*_. The dashed vertical line shows the theoretical optimal flee angle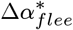, as by Hall et al. (11).

**Fig. S9.**
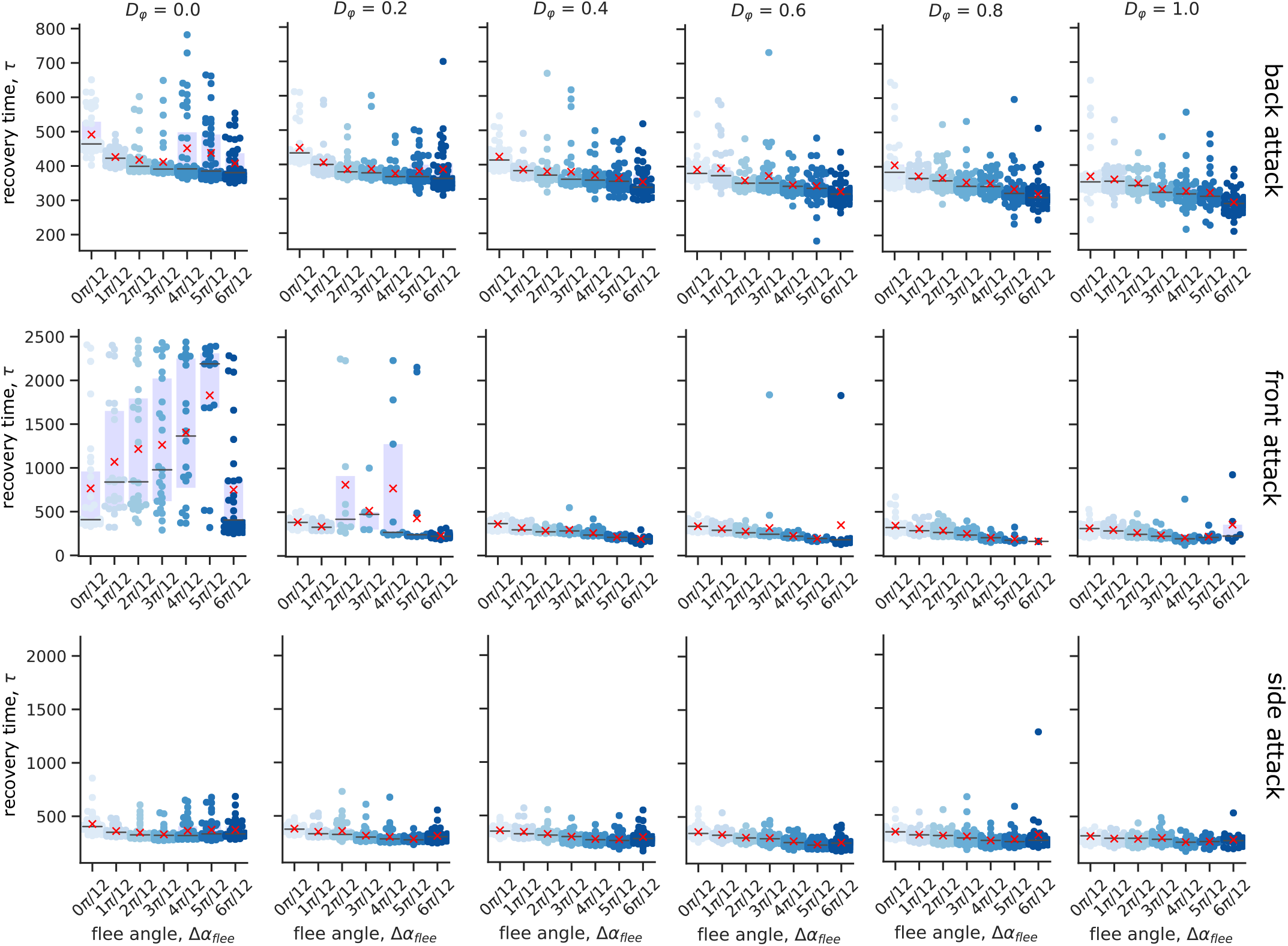
Collective prey recovery time *τ*, depending on the prey flee angle Δ*α*_*flee*_, prey orientational noise *D*_*φ*_ and the predator’s attack direction. The cross indicates the mean and the horizontal line above the boxplot indicates the median of the 40 simulation realizations for the corresponding parameter combination. The results show that the presence of the flee angle (Δ*α*_*flee*_ *>* 0*°*) allows for a faster collective recovery *τ* compared to Δ*α*_*flee*_ = 0*°*, particularly, after back and side attacks. The predator:prey speed ratio is set to 2 : 1.

**Fig. S10.**
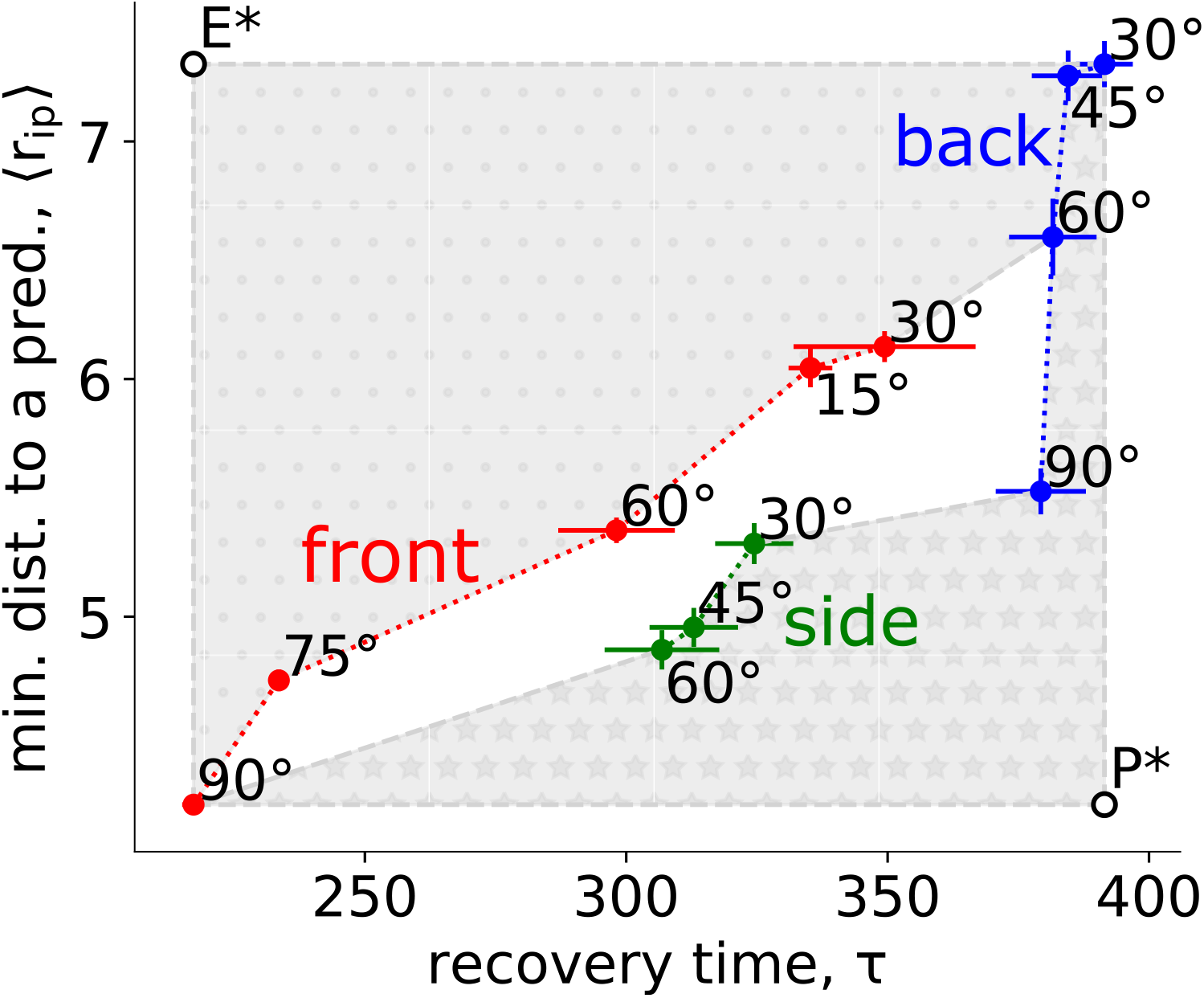
Pareto fronts of attack directions, conditioned on the Pareto optimal prey escape. *E*^*\**^ denotes the ideal solution for the prey across all attack directions as the maximum value of minimal averaged prey distance to a predator ⟨*r*_*i,p*_⟩ and the minimum value of collective recovery time *τ*. *P* ^*\**^ denotes the ideal solution for the predator with the opposite goals to the prey (i.e., minimise ⟨*r*_*i,p*_⟩ and maximise *τ*). The grey dotted area towards *E*^*\**^ depicts the infeasible region for the prey, with the border being the Pareto front of attack directions for the prey (front attack with Δ*α*_*flee*_ ∈ *{*90*°*, 75*°*, 60*°*, 15*°*, 30*°}*, back attack with Δ*α*_*flee*_ ∈ *{*30*°*, 45*°*, 60*°}*). The grey starred area towards *P* ^*\**^ depicts the infeasible region for the predator, with the border being the Pareto front of attack directions for the predator (front attack with Δ*α*_*flee*_ = 90*°*, side attack with Δ*α*_*flee*_ ∈ *{*60*°*, 45*°*, 30*°}*, and back attack with Δ*α*_*flee*_ ∈ *{*90*°*, 60*°*, 45*°*, 30*°}*). Notably, side attack irrespective of the prey flee angle does not lie on the Pareto front of prey attack directions.

## Notes

### Competing Interest Statement

The authors have declared no competing interest.

